# An Immune Cell Atlas Reveals Dynamic COVID-19 Specific Neutrophil Programming Amenable to Dexamethasone Therapy

**DOI:** 10.1101/2021.04.18.440366

**Authors:** Sarthak Sinha, Nicole L. Rosin, Rohit Arora, Elodie Labit, Arzina Jaffer, Leslie Cao, Raquel Farias, Angela P. Nguyen, Luiz G. N. de Almeida, Antoine Dufour, Amy Bromley, Braedon McDonald, Mark Gillrie, Marvin J. Fritzler, Bryan Yipp, Jeff Biernaskie

## Abstract

SARS-CoV-2 is a novel coronavirus that causes acute respiratory distress syndrome (ARDS), death and long-term sequelae. Innate immune cells are critical for host defense but are also the primary drivers of ARDS. The relationships between innate cellular responses in ARDS resulting from COVID-19 compared to other causes of ARDS, such as bacterial sepsis is unclear. Moreover, the beneficial effects of dexamethasone therapy during severe COVID-19 remain speculative, but understanding the mechanistic effects could improve evidence-based therapeutic interventions. To interrogate these relationships, we developed an scRNA-Seq and plasma proteomics atlas (biernaskielab.ca/COVID_neutrophil). We discovered that compared to bacterial ARDS, COVID-19 was associated with distinct neutrophil polarization characterized by either interferon (IFN) or prostaglandin (PG) active states. Neutrophils from bacterial ARDS had higher expression of antibacterial molecules such as PLAC8 and CD83. Dexamethasone therapy in COVID patients rapidly altered the IFN^active^ state, downregulated interferon responsive genes, and activated IL1R2^+ve^ neutrophils. Dexamethasone also induced the emergence of immature neutrophils expressing immunosuppressive molecules ARG1 and ANXA1, which were not present in healthy controls. Moreover, dexamethasone remodeled global cellular interactions by changing neutrophils from information receivers into information providers. Importantly, male patients had higher proportions of IFN^active^ neutrophils, a greater degree of steroid-induced immature neutrophil expansion, and increased mortality benefit compared to females in the dexamethasone era. Indeed, the highest proportion of IFN^active^ neutrophils was associated with mortality. These results define neutrophil states unique to COVID-19 when contextualized to other life-threatening infections, thereby enhancing the relevance of our findings at the bedside. Furthermore, the molecular benefits of dexamethasone therapy are also defined, and the identified pathways and plasma proteins can now be targeted to develop improved therapeutics.

## COVID-19 ARDS host responses contextualized to bacterial ARDS

A broad array of infections including SARS-CoV-2 and bacterial sepsis can induce acute respiratory distress syndrome (ARDS), respiratory failure and death^1–3^. Neutrophils are thought to be key drivers of both COVID-19 and bacterial ARDS^4–6^, yet it is unclear if this is related to intrinsic and/or irreversible cellular responses. While recent studies have leveraged single-cell transcriptomics to dissect peripheral^7–9^and bronchoalveolar fluid ^10–12^immune landscapes driving COVID-19 pathogenesis, the protocols used can inadvertently exclude the majority of polymorphonuclear granulocytes, including neutrophils, as they are highly sensitive cells with low RNA (and high RNase) content. Here, we employ whole-blood-preserving protocols that capture all major immune cell types from critically ill patients admitted to intensive care units (ICUs) (Extended Fig 1). All samples taken from COVID-19 patients were assessed by bacterial culture and tested negative. All COVID-19 patients tested positive by PCR for SARS-CoV-2, and we previously confirmed an absence of viral mRNA in any circulating immune cells in a subset of patients^13^. However, a plasma proteomic screen for SARS-CoV-2 specific viral proteins in all samples revealed detection of one or more viral proteins in COVID-19 patient serum (Extended Fig 2a). Furthermore, we compared patient samples from COVID-19 ARDS to bacterial sepsis with ARDS (herein referred to as bacterial ARDS) (Extended Fig 2b), as there were unusually low admissions to ICU with viral pneumonias/ARDS during the period studied, likely due to COVID-19 public health measures.

Patient cohorts had comparable ages, sex, days on life support and time in hospital, but COVID-19 patients had broader racial diversity (Extended Fig 2c,d, Extended Data Table 1). Bacterial ARDS induced significant neutrophilia, and relative thrombocytopenia compared to the near normal circulating neutrophil numbers in COVID-19, while both had similar degrees of lymphopenia (Extended Fig 2e). Both cohorts had comparable PaO2 / FiO2 (P/F) ratios, which is an indicator of the severity of ARDS^14^, but bacterial ARDS patients had significantly more kidney injury demonstrated by higher levels of serum creatinine (Extended Fig 2f). We further compared families of soluble inflammatory markers (Extended Fig 2g) used to distinguish prototypical states, including those identified during cytokine storm (Extended Fig 2h) and cytokine release syndrome (Extended Fig 2i)^15^, which demonstrated similar soluble cytokine and chemokine responses between the infections. Therefore, in the context of life-threatening bacterial ARDS, COVID-19 ARDS patients had normal neutrophil counts, comparable IL-6 levels, and less organ failure as indicated by serum creatinine levels, all of which have been previously proposed as markers of COVID disease severity^16,17^. This prompted us to further investigate immune states and composition in response to COVID-19 compared to bacterial ARDS.

The online companion atlas (biernaskielab.ca/COVID_neutrophil) contains accessible scRNAseq data performed on freshly obtained whole blood at timepoint 1 (t1, <72h after ICU admission) and at timepoint 2 (t2, 7 days after t1) (Fig 1a). Cellular identity was mapped to 30 immune cell types/states using a UMAP projection from 21 patients and 86,935 cells (Fig 1b, Extended Figure 3a). Global magnitude of gene expression was directly compared between COVID-19 and bacterial ARDS patients (Extended Data Table 3), which revealed a more globally altered distribution of differential expression at t1 than at t2. Altered regulation of genes was most pronounced in neutrophils at t1, with lower neutrophil gene expression in COVID-19 compared to bacterial ARDS (Fig 1c; Extended Fig 3b-c). At t2, the global alterations in gene expression when comparing COVID-19 to bacterial ARDS were most pronounced in plasmablasts (Fig 1d; Extended Fig 3d–e). We further compared and quantified the proportions of known peripheral blood cellular constituents, which highlighted significant differences in CD4 T cells, CD8 T cells and NK cells (Extended Fig 3f). These data highlight that significant global differences in immune cell gene expression exist between COVID-19 ARDS and bacterial ARDS.

**Figure 1.**
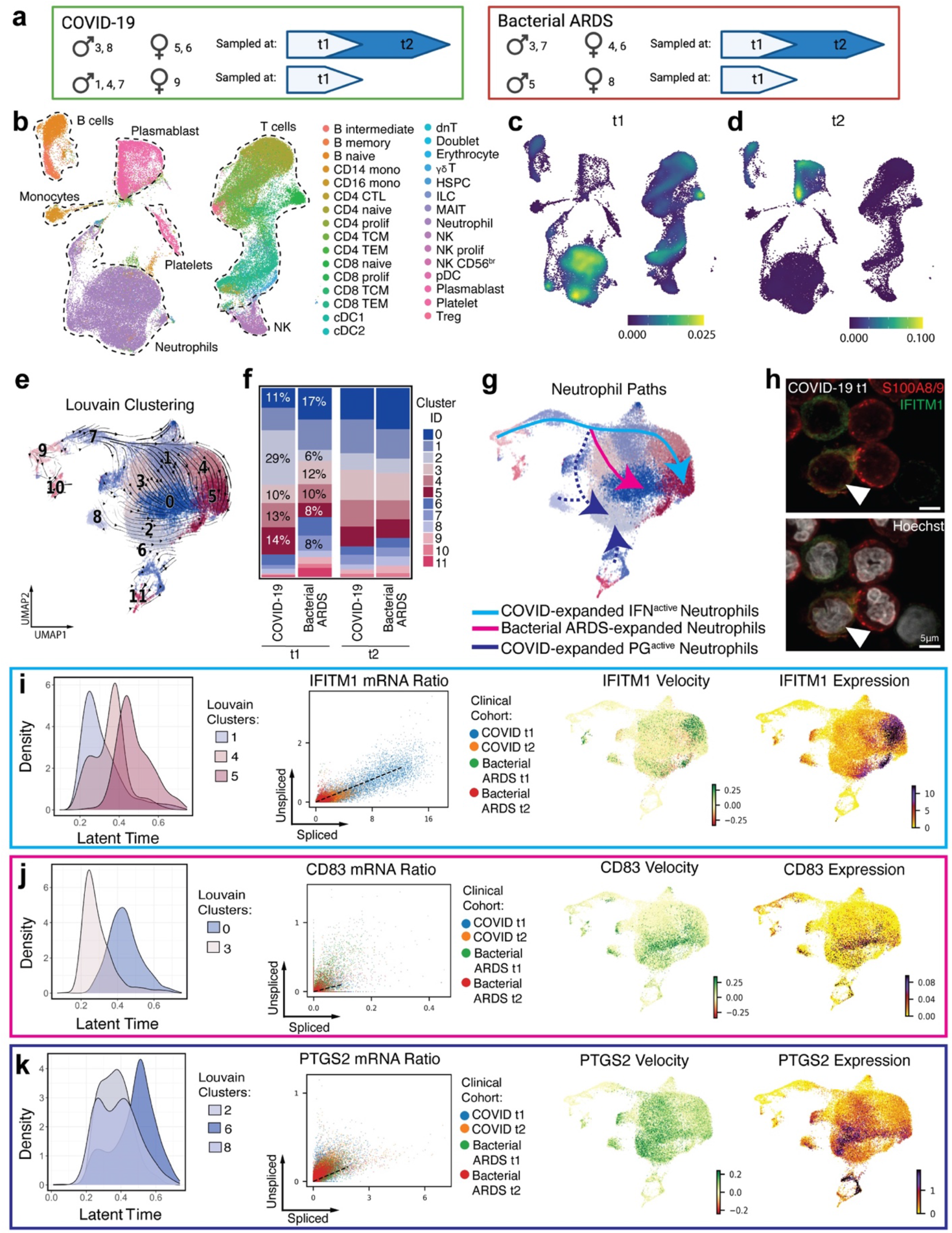
COVID-19 alters neutrophil maturation. **a.** Schematic summarizing patients with COVID-19 and bacterial sepsis profiled at t1 and t2. **b.** UMAP projection of 86,935 whole blood cells from 21 patient samples, coloured by Azimuth reference-mapped immune cell states. **c-d.** Kernel density estimates depicting magnitude of molecular response elicited by immune cell subsets during COVID-19 compared to Bacterial ARDS at t1 (c) and t2 (d) calculated by summing DEG fold changes for each cell state shown in Panel a. **e.** UMAP plotting RNA velocity analysis of 29,653 subclustered neutrophils undergoing state transitions, coloured by cluster ID. **f.** Stacked bar plot depicting cluster composition of clinical cohorts examined. **g.** UMAP coloured by neutrophil clusters and overlaid with summary path curves based on vector fields and neutrophil state compositions in Panel d and e, respectively to determine neutrophil states. **h.** Immunocytochemistry for S100A8/A9 (red) and IFITM1 (green) expression on leukocyte-rich preparation from COVID-19 donor at tl. **i-k.** Transcriptional kinetics driving expansion of IFN^active^ (i), Bacterial ARDS-enriched (j), and PG^active^ (k) neutrophils. Latent time distribution of trajectory-associated louvain clusters (left), phase portraits with equilibrium slopes of spliced–unspliced ratios (center), and RNA velocity and gene expression (right) of selected genes driving divergent maturation trajectories. Phase portraits are coloured by clinical cohort.

## COVID-19 drives specific neutrophil maturation states

Neutrophils are a primary participant in the development of ARDS^18^; yet despite similar severity of ARDS between our bacterial and COVID-19 cohorts, the numbers of circulating neutrophils from clinical cell counts were significantly different (Extended Fig 2d). We hypothesized that neutrophil qualitative states may be important determinants of disease. Neutrophils were subjected to velocity analysis^19,20^ to reconstruct maturation dynamics. Louvain clusters (Fig 1e), clinical cohort, individual patient, and velocity length were overlayed on velocity vector fields (Extended Fig 4a–d). The proportions of distinct neutrophil states were compared at t1 and this revealed a divergent expansion of IFN^active^ neutrophils (clusters 2, 4 and 5) marked by IFITM1 expression in COVID-19, which became similar to bacterial ARDS at t7 (Fig 1f–h). Expression of IFITM1 in neutrophils from COVID-19 patients at t1 was confirmed by immunofluorescent staining for IFITM1, colocalized with S100A8/9 and typical neutrophil nuclear morphology. Relative to healthy donors, the IFN^active^ population in both COVID-19 and bacterial ARDS patients were elevated (Extended Fig 4h–k), suggesting that infections dramatically alter neutrophil dynamics and that comparing COVID-19 neutrophils to healthy neutrophils may only reveal broad features separating pathogen-challenged versus non-challenged (homeostatic) neutrophils. Hence, to map pathogen-activated neutrophils dynamics with high resolution, subsequent analyses employ principal components with top loading genes that distinguish different pathogen-activated states arising during COVID-19 and bacterial sepsis for downstream dimensionality reduction.

Classically, peripheral neutrophils are considered terminally differentiated and non-dividing, however the increase in velocity length suggested the ability to alter phenotypic states once in circulation along specific paths or ‘lineages’. COVID-19 neutrophils followed unique maturation paths compared to bacterial ARDS, culminating in three distinct terminal states: Interferon active (IFN^active^), prostaglandin active (PG^active^) or bacterial ARDS enriched (Fig 1e–g; Extended Fig 4e). Interestingly, the apex of this trajectory was marked by high velocity lengths, characteristic of cells undergoing differentiation (Extended Fig 4c, d). COVID-19 neutrophils preferentially transitioned from the apex of the trajectory, which was an immature state (TOP2A expressing; Extended Fig 4e) to an IFN responsive state characterized by IFITM1, IFITM2 and IFI6 expression (Cluster 1 to 4 and 5; Fig 1i; Online Atlas). This is clearly illustrated in Extended Video 1. This immature state was not present in healthy controls, though it is present in both comparator groups, suggesting these states are liberated into circulation upon pathogen exposure (Extended Fig 4h–k). The lineage relationship was less clear for COVID-19 enriched PG^active^ clusters defined by prostaglandin responsive genes (clusters 2, 6 and 8), with notable increases in PTGER4 and PTGS2 (or COX2), a proposed therapeutic target in COVID-19^21^ (Fig 1k; Extended Fig 4f, g, Online Atlas). The dominant conventional bacterial ARDS state was characterized by antibacterial proteins CD83^22^, CD177, and PLAC8^23^ (cluster 3 to 0; Fig 1j; Online Atlas). Taken together, this data demonstrated that peripheral neutrophils have dynamic programming abilities which result in COVID-19 specific neutrophil polarization defined by the emergence of IFN^active^ and PG^active^ neutrophil states.

## Unique transcriptional regulatory pathways drive neutrophil maturation in COVID-19

Rapid and robust IFN responses protect against COVID-19 severe disease, while delayed responses could exacerbate systemic and pulmonary inflammation^24,25^. Moreover, neutrophil IFN responses are not traditionally considered during infections and neutrophils are generally considered to be homogenous, with a uniform proinflammatory capacity. Global neutrophil expression aligned with neutrophil state specific markers, such as interferon response genes (IFITM1, RSAD2, IFI6, and ISG10), being more highly expressed in COVID-19 neutrophils (Fig 2a; Extended Fig 4f). The inverse was the case for anti-bacterial proteins like PLAC8 (Fig 2a; Online Atlas). However, the discovery of differential neutrophil states prompted further exploration of the factors driving neutrophil state polarization. Gene regulatory network reconstruction using SCENIC analysis^26^ revealed differentially activated transcription factors STAT1, IRF2 and PRDM1 in COVID-19 (Fig 2b), while bacterial ARDS neutrophils had increased prototypical granulocyte transcription factors such as CEBPA, CEBPB, STAT5B and less defined factors such as NFE2 (Fig 2b, Online Atlas). PRDM1 activation was most pronounced in the IFN^active^ neutrophil population and was likely responsible for driving expression of interferon response elements (IFIT1, ISG15, IFI6) and antiviral signaling, such as RSAD2 and STAT1 (Fig 2c; Online Atlas). A hallmark of PG^active^ neutrophil polarization was the activation of an E2F4 pathway (Fig 2d), while neutrophil programming during bacterial ARDS included STAT5B (Fig 2e). To summarize, in response to COVID-19, neutrophils were polarized by unique transcriptional regulation towards one of two main populations, either an IFN^active^ population or a PG^active^ population (Fig 2f).

**Figure 2.**
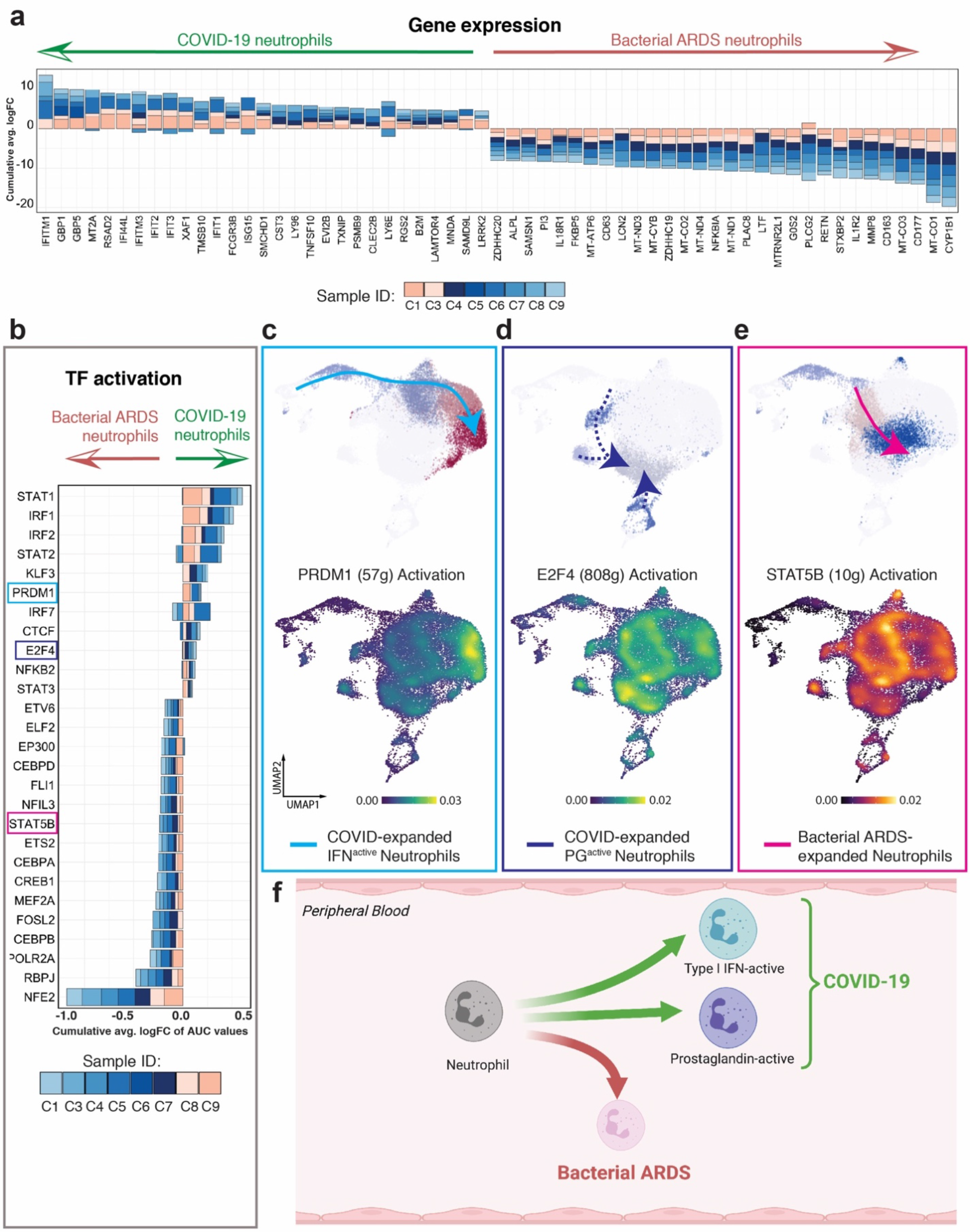
Distinct regulatory programs drive divergent neutrophil maturation. **a.** Consensus neutrophil DEGs upregulated (positive FC) or suppressed (negative FC) during COVID-19 in at least 3 of 8 patients at t1 relative to Bacterial ARDS. **b.** Differentially activated consensus transcription factors (TFs) in neutrophils from patients with COVID-19 relative to bacterial ARDS at t1. Stacked bars depict logFC contributions of each COVID-19 patient. **c-e.** Gene-regulatory networks preferentially driving IFN^active^ (PRDM1, c), PG^active^ (E2F4, d), and bacterial ARDS-enriched (STAT5B, e) neutrophil states. Scale bars depict kernel density estimates approximating magnitude of TF activation inferred by SCENIC-calculated AUCell scores. **f.** Schematic summarizing neutrophil fates favoured during COVID-19 versus bacterial ARDS.

## Dexamethasone alters immune cell dynamics and plasma proteomic milieu

Conventional therapeutics have limited efficacy for COVID-19, and while dexamethasone offers a moderate benefit, the RECOVERY trial reported the benefit was greatest in the most severely affected patients^27^. However, the mechanisms underlying this benefit are unclear and not universal, so opportunity exists to optimize or better target this therapy. In our cohort, median time between dexamethasone administration to t1 blood draw (within 72 hours of ICU admission) was 31 hours (Fig 3a, Extended Figure 5a, Extended Table 1). Global differences in transcription were apparent at t1 with clear upregulation of genes in neutrophils and some T cell subsets in COVID-19 patients treated with dexamethasone versus those that were not treated (Fig 3b–d, Extended Figure 5b, Extended Data Table 4). Dexamethasone globally downregulated genes at t1, including in naïve B cells, plasmablasts and some T cells (Extended Figure 5b-d). At t2 gene upregulation occurred in adaptive immune cells, including naïve and effector CD8 T cells, with limited alterations in the innate myeloid cell lineages including neutrophils. However, neutrophils demonstrated clear down regulation of genes at t2, as did CD4 naïve and central memory T cells (Extended Figure 5e, f). Proportionally, at t1, dexamethasone administration was associated with an increase in cytotoxic CD4 T cells, naïve B cells, plasmablasts, and decreased proliferating NK cells, and CD4 effector memory cells (Extended Fig 5g). By t2, dexamethasone was associated with suppressed neutrophil proportions in circulation compared to untreated controls (13% vs 41%, Extended Fig 5g). Plasma proteomics from the same cohort revealed that dexamethasone suppressed 10 host proteins (S100A8, S100A9, SERPINA1, SERPINA3, ORM1, LBP, VWF, PIGR, AZGP1, CRP) that others have previously identified as biomarkers distinguishing severe COVID-19 cases from mild to moderate counterparts (full host proteome quarriable via Online Atlas; Extended Table 2)^28–31^. Suppression of calprotectin (S100A8/S100A9) and neutrophil serine proteases (SERPINA1 and SERPINA3), paired with depletion of neutrophil proportions, implicates the modulation of neutrophil-related inflammatory processes as a method of action for dexamethasone treatment.

**Figure 3.**
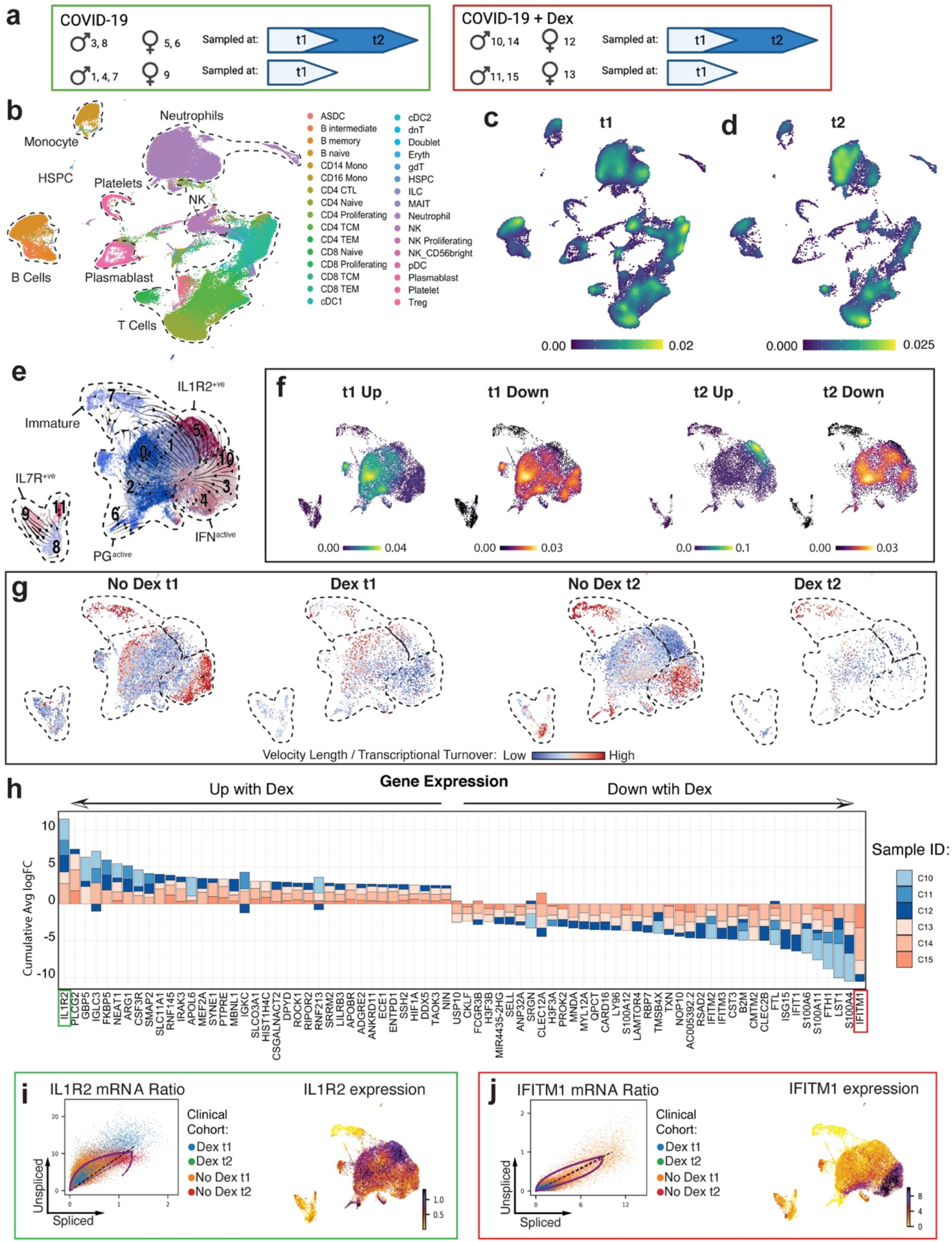
Dexamethasone suppresses IFN programs and depletes IFN^active^ neutrophils in COVID-19. **a.** Schematic summarizing COVID-19 patients treated with or without dexamethasone profiled at t1 and t2. **b.** UMAP projection of 80,994 whole blood cells from 21 patient samples, coloured by Azimuth reference-mapped immune cell states. **c-d.** Kernel density estimates depicting magnitude of molecular response elicited by immune cell subsets following Dexamethasone treatment t1 (c) and t2 (d) calculated by summing DEG fold changes for each cell state shown in Panel A. **e.** Neutrophil states overlaid on a UMAP of 23,193 subclustered neutrophils from Dexamethasone- and non-Dexamethasone-treated COVID-19 patients, colored by cluster ID. **f.** Magnitude of molecular response elicited by each neutrophil state post-Dexamethasone treatment calculated by summing DEG fold changes for each cell state shown in Panel d. **g.** RNA velocity vector length (indicating rate of differentiation/state transition) in Dexamethasone- and non-Dexamethasone-treated neutrophils at t1 and t2. **h.** Consensus neutrophil DEGs upregulated (positive FC) or suppressed (negative FC) post-Dexamethasone in at least 3 of 6 COVID-19 patients at t1 relative to non-Dexamethasone COVID-19 controls. Stacked bars depict logFC contribution of each Dexamethasone-treated patient. **i-j.** Differential splicing kinetics drives activation of IL1R2 (i) and suppression of IFITM1 expression (j) post-Dexamethasone treatment. Phase portraits show equilibrium slopes of spliced–unspliced mRNA ratios. Green denotes most upregulated and red denotes most down regulated differentially expressed genes with COVID-19 (f).

## Dexamethasone therapy restrains neutrophil IFN programs

Due to the early and sustained effects of dexamethasone on gene expression in neutrophils, the effects of dexamethasone therapy on neutrophil functional states were evaluated. Neutrophil reclustering again identified immature neutrophils at the apex of the maturation trajectory, accelerating and exhibiting maximal divergence prior to PG^active^ and IFN^active^ state commitments (Fig. 3 d, Extended Fig 6a-e). Interestingly, we also identified IL7R^+ve^ neutrophils (comprising roughly 8% of total neutrophils) whose trajectories remained completely separate (Fig. 3 d, Extended Fig 6g, j) suggesting an entirely distinct neutrophil state. Initially, dexamethasone was associated with increased global transcription in PG^active^ neutrophils, while ongoing therapy resulted in the emergence of a PG^active^ neutrophils concomitant with high IL1R2 expression (IL1R2^+ve^) (Fig. 3 e). Conversely, dexamethasone had a pronounced attenuation of global transcription of IFN^active^ neutrophils at t1 and t2 (Fig 3 e, f). Remarkably, dexamethasone administration at t1 halted dynamic state changes in IFN^active^ and IL7R^+ve^ neutrophils, followed by preferential depletion of IFN^active^ subsets (Fig 3 g). Indeed, dexamethasone was associated with a reduction in IFN^active^ neutrophils to a proportion more similar to that detected in healthy controls (9% post-Dex at t2 versus 10% in healthy controls) (Fig. 4a, Extended Fig 4h-k). Although collection of airway samples (i.e. bronchoalveolar lavage fluid; BALF) was not feasible at our institution, we leveraged two recent BALF scRNA-Seq datasets^11,32^ to assess whether IFN^active^ neutrophils dominate the bronchoalveolar landscape during severe COVID-19. Projection of CSF3R^+^S100A8^+^S100A9^+^ BALF neutrophils onto our reference revealed: a. 1.5 FC expansion of IFN^active^ neutrophils in severe COVID-19 relative to moderate disease (77% vs 52%, Extended Fig 7a-b), b. preferential activation of IFN-stimulated genes such as IFITM1, IFITM2, IFI6, IRF7, and ISG20 in severe COVID-19 neutrophils (Extended Fig 7c), and c. 4.7 FC greater IFN^active^ neutrophils in COVID-19 relative to bacterial pneumonia patients (14% vs 3%, Extended Fig 7d-f). Albeit anecdotal, in our whole blood cohort, the IFN^active^ neutrophil state was dominant in patient S7 ^32^, an 80-year-old male with remarkably high viral titers who succumbed to COVID-19 complications within 3-4 days of sampling (Extended Fig 7f). Consensus DEG analysis highlighted that upregulation of IL1R2, a decoy receptor that sequesters IL-1, and downregulation of IFITM1 were the most prominent discriminating features of treatment with steroids (Fig. 3h). Additionally, dexamethasone attenuated neutrophil expression of IFN pathways more broadly, including the reduction of IFITM1-3, IFIT1, ISG15 and RSAD2 (Fig 3h). Examination of unspliced pre-mRNA to mature spliced mRNA ratios supported the notion that induction of immunoregulatory systems (i.e., IL-1R2; Fig 3 i) and suppression of IFN (i.e., IFITM1; Fig 3 j) programs were driven by differential splicing kinetics.

## Dexamethasone therapy intensifies neutrophil immunosuppressive function

Corticosteroid therapy shifted neutrophil state compositions. While IFN^active^ neutrophils were significantly depleted by seven days of therapy, there was >2-fold expansion in immature neutrophils relative to non-treated COVID-19 controls (Fig 4a; Extended Fig 6 h, i), which were absent in the healthy controls. Albeit anecdotal, the dominance of IFN^active^ neutrophils at t1 in the patient who succumbed to COVID-19 in the non-dexamethasone cohort further supports depletion of IFN^active^ neutrophils as a mechanism by which dexamethasone is protective (Extended Fig 8 g-j). Assessment of gene regulatory networks demonstrated that IRF7 and MEF2A exhibited opposing activation patterns, with IRF7 being the most suppressed and MEF2A the most enhanced transcription factors identified with dexamethasone, which correlates with the emergence of PG^active^ and IL1R2^+ve^ states and attenuation of the IFN^active^ neutrophil states (Fig 4b, Extended Fig 6k-m). To assess the generalizability of the dexamethasone regulated DEGs identified in our cohort, we asked whether they accurately predicted mortality due to COVID-19 in a larger validation cohort. By leveraging a whole blood bulk RNA-Seq dataset from 103 COVID-19 patients^33, 34^, we scored each sample by the aggregated expression of dexamethasone suppressed DEGs at t1 and t2 (Extended Data Table 3). Interestingly, suppressed DEGs at t2 (but not t1) proved to be a far superior predictor of 28-day mortality (AUC: 0.78, CI: 0.67 −0.89) compared to clinical severity scales such as sequential organ failure assessment (SOFA) (AUC: 0.67, CI: 0.51-0.82) across all classification thresholds (Fig 4c).

**Figure 4.**
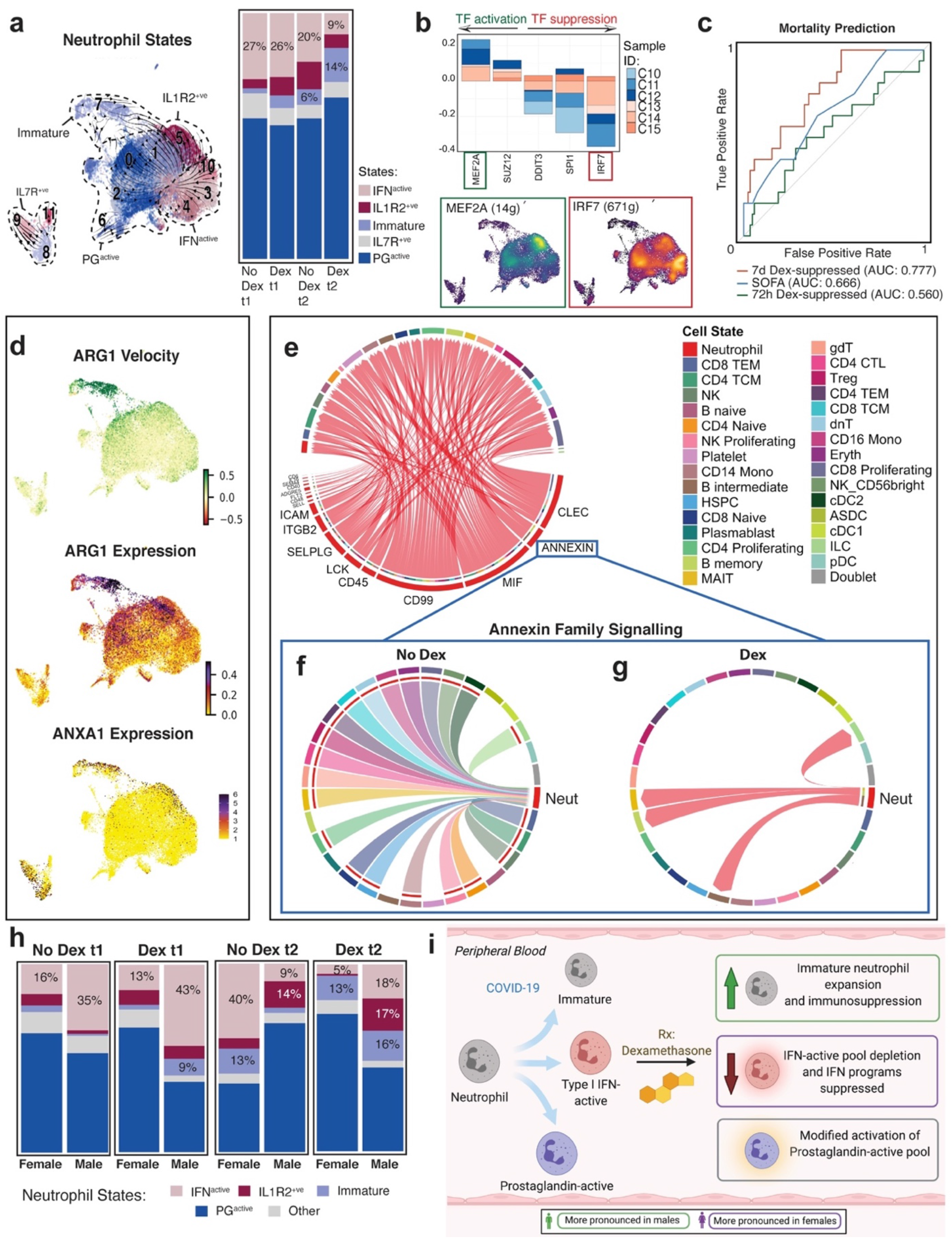
Dexamethasone expands immunosuppressive neutrophils and their interactions in COVID-19. **a.** Neutrophil states mapped onto Louvain-clustered UMAP, with comparison of neutrophil composition between dexamethasone- and non-dexamethasone-treated samples at t1 and t2. **b.** Consensus TFs activated or suppressed post-dexamethasone in at least 3 of 6 patients at t1 and predicted activity of MEF2A and IRF7, two of the most differentially regulated TFs post-dexamethasone. **c.** Receiver operating characteristic (ROC) curves assessing the discriminatory capacity of dexamethasone suppressed DEGs at t1, t2, and sequential organ failure assessment (SOFA) scores for predicting 28-day mortality in a validation cohort of 103 bulk whole blood RNA-Seq samples where 17 cases were fatal. **d.** Immature and IL1R2^+ve^ neutrophil subsets express high levels of immunosuppressive neutrophil marker ARG1 and ANXA1. **e.** Neutrophil-driven signaling pathways induced post-dexamethasone, identified using CellChat (MHC-I signalling filtered out). **f**, **g**. Topology of annexin signalling without (e) and with dexamethasone (f) treatment (edges filtered to those where neutrophils function as senders or recipients of annexin signals). **h.** Neutrophil state composition separated by sex and dexamethasone status at t1 and t2. **i.** Schematic summarizing the effects of dexamethasone on neutrophil fates and function in COVID-19 following dexamethasone treatment.

Unexpectedly, steroid administration was associated with an increase in circulating immature neutrophils, which highly expressed TOP2A, and activated ATF4 and JDP2, transcription factors seen in undifferentiated cells or those undergoing nuclear reprogramming (Extended Fig 6h). Interestingly, these immature neutrophils expressed high levels of ARG1, ANXA1 (Fig 4d), and CD24 (both mRNA and protein; Extended Fig 6 i), also suggesting an immunomodulatory role^35,36,37–39^ that was expanded with dexamethasone treatment. Both ARG1 and ANXA1 express glucocorticoid response elements, supporting direct regulation by dexamethasone treatment^40,41^.

To further understand the role of neutrophils during COVID-19 and the effects of dexamethasone, we investigated cellular connectomes. Cellular interactions between many cell types (including highly interactive neutrophils) were noted (Extended Fig 8a), and dexamethasone altered the globally predicted interactions by suppressing intercellular signalling, in both number and strength of interactions (Extended Fig 8b, c). Dexamethasone enhanced (Fig 4e) and suppressed (Extended Fig 8d) a number of unique neutrophil-driven signalling networks. Of note, annexin family signalling, which was enhanced in the immature neutrophils and represent powerful immunomodulators, were augmented between neutrophils and the other circulating immune cells when patients received dexamethasone (Fig 4e). Of note is the direction of annexin family signaling, which switched from incoming toward neutrophils without dexamethasone treatment to being almost entirely outgoing from neutrophils toward B intermediate and memory cells and MAIT cells following dexamethasone (Fig 4f, g, Extended Fig 8e, f). Therefore, dexamethasone directly altered neutrophil functional states, by promoting expansion of an ARG1+/ANXA1+ immature state with immunosuppressive features and altered the global communication structure such that neutrophils became active instructors of some peripheral immune cells.

## Neutrophil response to dexamethasone is sexually dimorphic

Given the apparent clinical benefit of dexamethasone is more evident in males^27^, and since males are predisposed to more severe COVID-19 presentations and outcomes^42^, we surmised that dexamethasone incites sexually dimorphic immunosuppressive effects. Our retrospective province-wide audit comparing 72 pre-dexamethasone (51 M, 21 F) versus 1,581 post-dexamethasone (1013 M, 568 F) treated ICU-admitted patients confirmed a preferential mortality benefit in male COVID-19 patients (Extended Fig 9a, b). While dexamethasone modulated 525 neutrophil DEGs across both sexes, while 892 were uniquely modulated in either males or females (Extended Data Table 5). Amongst the jointly modulated DEGs, a subset (24 of 525) exhibited statistically significant dimorphism in either magnitude or direction of regulation (Extended Fig 9c, d). Interestingly, while neutrophils were depleted in both sexes post-dexamethasone, this was particularly pronounced in males (1.9 FC higher in males at t1 and 3.4 FC higher in males at t2, Extended Fig 9e). Of the two salient neutrophil state alterations, an immature (ARG1^+ve^ immunosuppressive) state was preferentially expanded with dexamethasone in males (Extended Fig. 9e), whereas ISGs were preferentially suppressed (Extended Fig. 9f) and IFN^active^ states were depleted in females (Extended Fig. 9g-h) at both t1 and t2 (Fig 4h, i). Sexually dimorphic effects of dexamethasone on neutrophil maturation kinetics may in part explain these state alterations. Dynamoreconstructed vector dynamics revealed that dexamethasone slowed IFN^active^ transitions (Extended Fig. 9i) whilst accelerating immature (ARG1^+ve^ immunosuppressive) neutrophil differentiation in females (Extended Fig. 9j) ultimately leading to a diminished immature neutrophil progenitor pool.

## Conclusions

Surviving SARS-CoV-2 infection depends on striking a temporal balance between inciting viral clearance immune programs during the early stage and subsequently restraining those same programs at later stages to limit immunity-induced tissue damage. IFN signaling stands at the nexus between antiviral immunity and over active effector immune programs that inadvertently compromise tissue function and threaten survival^43^. Our work uncovered downstream IFN signalling as a signature of a stable neutrophil state that is selectively expanded during late stage COVID-19 infection from a common pool of immature progenitors. Given that inborn errors ^25^ and suppressed *early stage ^6^* IFN signalling predicts COVID-19 severity, increased IFN^active^ neutrophils in females correlated with decreased mortality^44^, and early initiation of IFN therapy has been suggested to mitigate disease severity ^45,46^, one may posit that IFN activity in neutrophils represents a concerted host antiviral program.

Interestingly, immunosuppression with dexamethasone, a corticosteroid known to improve mortality in hospitalized COVID-19 patients^27^, was associated with suppressed COVID19-specific IFN regulatory networks and depleted COVID19-enriched IFN^active^ neutrophils in favour of expanding immature (ARG1^+^ immunosuppressive) neutrophils. These altered neutrophil states shared striking resemblances to bacterial ARDS, suggesting installation of generalized microbicidal programs ameliorate the overzealous neutrophil responses during COVID-19 (and perhaps during other viral infections). While neutrophil ISG activation may promote anti-viral immunity during early stages of SARS-CoV-2 infection, sustained IFN activation during *late stages* (e.g., critically ill patients requiring intensive care) could drive immunopathology of COVID-19. Indeed, positive correlation between neutrophil Type 1 IFN programs and COVID-19 severity^7,47^ paired with our observation that IFN^active^ neutrophils dominate the bronchoalveolar microenvironment during severe COVID-19 ^11^directly support this view. Immunotherapies that support the innate antiviral immune response by decoupling IFN-exaggerated neutrophil response whilst reinforcing acquisition of suppressor states may limit the pathogenic potential of neutrophils and provide tremendous clinical benefit for treating severe COVID-19.

There are three major limitations of our study. First, <u>non-random group allocation (since the timing of the RECOVERY trial made dexamethasone standard of care overnight) and small sample size may inadvertently introduce selection bias and limit generalizability of dexamethasone findings</u>. Second, comparisons were against bacterial ARDS, and not related respiratory viral infections (i.e., H1N1 influenza) since public health measures eradicated such cases; this precludes assessment of whether the dynamics defined are specific to SARS-CoV-2. Finally, a subset of patients sampled at t1 were discharged from ICU prior to t2 collection (non-random or non-ignorable missing data), precluding unbiased estimation of temporal changes between timepoints.

## Methods

### Patient enrolment

All patients were enrolled following admission to any of the four adult intensive care units at South Health Campus, Rockyview General Hospital, Foothills Medical Center or Peter Lougheed Center in Calgary, Alberta, Canada (Extended Fig 1). Patient admission to the ICU was determined by the attending ICU physician based on the need for life sustaining interventions, monitoring and life-support. The research teams did not participate in clinical decisions. Study inclusion required a minimal age of 18, the ability to provide consent, or for most participants, the ability of a surrogate decision maker to provide regained capacity consent. All participants required an arterial catheter for blood draws, but the insertion of this catheter was at the discretion of the attending medical team. Participants required a positive clinical RNA COVID-19 test prior to enrolment, and evidence of bilateral lung infiltrates and hypoxemia consistent with ARDS. At the time of sample collections, all COVID-19^+^ enrolled individuals were culture negative for concurrent bacterial infections in the blood, urine, and sputum. The bacterial ARDS cohort required a negative COVID-19 test and a definitive microbiological diagnosis of bacterial pneumonia with chest imaging consistent with a diagnosis of ARDS. Patients were excluded from our study if they: 1. were on immunosuppressive therapies, 2. had established autoimmune disease, or 3. had active malignancy. Since tocilizumab or other immunomodulatory agents were not approved for use in patients with severe COVID-19 in Alberta over the timespan of this study, none of them received these medications. While bacterial sepsis patients received appropriate antibiotic treatments, none were prescribed immunosuppressive or steroid therapy. All bacterial sepsis patients had lung infections caused by gram-positive cocci (4 *Staphylococcus aureus* and 2 *Streptococcus pneumoniae*). Participants were required to have a definitive diagnosis and appropriate consent and samples collected within 72hrs of admission to the ICU in order to be included. Timepoint 1 (T1) refers to the first blood draw, while T2 was a repeat blood draw taken 7 days after T1, if the participant remained in the ICU, and had an arterial catheter. For each participant, whole blood was collected via the arterial catheter and immediately processed for analysis. Healthy blood donors were recruited by university-wide advertisement and required that participants were: 1. not on immunomodulatory medications, 2. were asymptomatic for SARS-CoV-2, 3. did not receive vaccination against SARS-CoV-2, and 4. did not have underlying immune disorders.

### Epidemiological analysis

We used the Alberta provincial eCRITICAL oracle-based analytics database (Tracer) to query and extract Alberta COVID-19 ICU cases and volumes for this study^48^. Aggregate data from sixteen individual adult ICUs was obtained over the study periods. The administration of dexamethasone was not possible to capture at an aggregate level; therefore, we queried the database for patients admitted to ICU prior to dexamethasone becoming standard of care in our Province (pre-dexamethasone era; January 2020 till May 31^st^, 2020) versus dexamethasone as standard of care for severe COVID-19 (June 1^st^, 2020, till May 31^st^, 2021). Tocilizumab was approved for use in Alberta March 11 2021, and a small supply (150 doses) was obtained for severe COVID-19 patients after this date.

### Human Study Ethics

All work with humans was approved by the Conjoint Health Research Ethics Board (CHREB) at the University of Calgary (Ethics ID: REB20-0481) and is consistent with the Declaration of Helsinki.

### Serum cytokine assessment

Cytokines, chemokines and soluble cytokine receptors were quantitated on multiplex arrays that included a 65 MIlliPLEX cytokine/chemokine (6Ckine, BCA-1, CTACK, EGF, ENA-78, Eotaxin, Eotaxin-2, Eotaxin-3, FGF-2, Flt-3L, Fractalkine, G-CSF, GM-CSF, GRO, I-309, IFNα2, IFNγ, IL-1α, IL-1β, IL-1ra, IL-2, IL-3, IL-4, IL-5, IL-6, IL-7, IL-8, IL-9, IL-10, IL-12 (p40), IL-12 (p70), IL-13, IL-15, IL-16, IL-17A, IL-18, IL-20, IL-21, IL-23, IL-28a, IL-33, IP-10, LIF, MCP-1, MCP-2, MCP-3, MCP-4, MDC, MIP-1α, MIP-1β, MIP-1d, PDGF-AA, PDGF-AB/BB, RANTES, SDF-1 a+b, sCD40L, SCF,TARC, TGFa, TNFa, TNFb, TPO, TRAIL, TSLP, VEGF) and a 14 MilliPLEX soluble cytokine (sCD30, sEGFR, sgp130, sIL-1RI, sIL-1RII, sIL-2Ra, sIL-4R, sIL-6R, sRAGE, sTNF RI, sTNF RII, sVEGF R1, sVEGF R2, sVEGF R3) arrays (Millipore Sigma, Oakville, ON, Canada) on a Luminex Model 200 Luminometer (Luminex Corporation, Austin, TX). EDTA-plasma samples were collected from each patient by venipuncture following a standard operating protocol (SOP) and stored at − 80C until tested. Each run included a full range of calibrators. The Mann-Whitney U test was used to compare groups and p-values were adjusted for multiple comparisons using Holm-Šídák stepdown method with alpha set to 0.05.

## Shotgun proteomics using Liquid Chromatography and Mass Spectrometry (LC-MS/MS)

The serum of COVID-19 patients (COVID-19 = 9, dexamethasone-treated = 4) and bacterial ARDS controls (N = 6) were collected and subjected to quantitative proteomics. The total protein concentrations were determined by Pierce™ BCA Protein Assay Kit (23225, ThermoFisher). A trichloroacetic acid (TCA)/acetone protocol was used to pellet 100μg of proteins per sample. Samples were subjected to a quantitative proteomics workflow as per supplier (Thermo Fisher) recommendations. Samples were reduced in 200mM tris(2-carboxyethyl)phosphine (TCEP), for 1h at 55°C, reduced cysteines were alkylated by incubation with iodoacetamide solution (50mM) for 20min at room temperature. Samples were precipitated by acetone/methanol, and 600μL ice-cold acetone was added followed by incubation at −20°C overnight. A protein pellet was obtained by centrifugation (8,000*g*, 10min, 4°C) followed by acetone drying (2min). Precipitated pellet was resuspended in100 μL of 50mM triethylammonium bicarbonate (TEAB) buffer followed by tryptase digestion (5μg trypsin per 100μg of protein) overnight at 37°C. TMT-6plex™ Isobaric Labeling Reagents (90061, Thermo Fisher) were resuspended in anhydrous acetonitrile and added to each sample (41μL TMT-6plex™ per 100μL sample) and incubated at room temperature for 1h. The TMT labeling reaction was quenched by 2.5% hydroxylamine for 15min at room temperature. TMT labeled samples were combined and acidified in 100% trifluoroacetic acid to pH < 3.0 and subjected to C18 chromatography (Sep-Pak) according to manufacturer recommendations. Samples were stored at −80°C before lyophilization, followed by resuspension in 1% formic acid before liquid chromatography and tandem mass spectrometry analysis.

Tryptic peptides were analyzed on an Orbitrap Fusion Lumos Tribrid mass spectrometer (Thermo Scientific) operated with Xcalibur (version 4.0.21.10) and coupled to a Thermo Scientific Easy-nLC (nanoflow liquid chromatography) 1200 System. Tryptic peptides (2μg) were loaded onto a C18 trap (75μm × 2cm; Acclaim PepMap 100, P/N 164946; ThermoFisher) at a flow rate of 2μL/min of solvent A (0.1% formic acid in LC-MS grade H2O). Peptides were eluted using a 120min gradient from 5 to 40% (5% to 28% in 105min followed by an increase to 40% B in 15min) of solvent B (0.1% formic acid in 80% LC-MS grade acetonitrile) at a flow rate of 0.3μL/min and separated on a C18 analytical column (75μm × 50cm; PepMap RSLC C18; P/N ES803A; ThermoScientific). Peptides were then electrosprayed using 2.1kV voltage into the ion transfer tube (300°C) of the Orbitrap Lumos operating in positive mode. For LC-MS/MS measurements with the FAIMS Pro (Thermo Fisher Scientific), multiple compensation voltages (CV) were applied, −40V, −60V, and −80V with a cycle time of 1 second. FAIMS was used to generate technical replicates from plex 1 to 6. The Orbitrap first performed a full MS scan at a resolution of 120,000 FWHM to detect the precursor ion having a m/z between 375 and 1,575 and a +2 to +4 charge. The Orbitrap AGC (Auto Gain Control) and the maximum injection time were set at 4 × 10^5^ and 50ms, respectively. The Orbitrap was operated using the top speed mode with a 3 second cycle time for precursor selection. The most intense precursor ions presenting a peptidic isotopic profile and having an intensity threshold of at least 2 × 10^4^ were isolated using the quadrupole (Isolation window (m/z) of 0.7) and fragmented using HCD (38% collision energy) in the ion routing multipole. The fragment ions (MS2) were analyzed in the Orbitrap at a resolution of 15,000. The AGC and the maximum injection time were set at 1 × 10^5^ and 105ms, respectively. The first mass for the MS2 was set at 100 to acquire the TMT reporter ions. Dynamic exclusion was enabled for 45 seconds to avoid of the acquisition of same precursor ion having a similar m/z (plus or minus 10ppm).

## Proteomic data and bioinformatics analysis

Spectral data acquired from the mass spectrometer were matched to peptide sequences using MaxQuant software (v.1.6.14)^49^. Due to a lack of direct compatibility with Maxquant, spectra generated using the FAIMS pro was first converted to MzXML using the FAIMS MzXML Generator from the Coon’s lab (https://github.com/coongroup/FAIMS-MzXML-Generator). Next, peptide sequences from the human proteome and Sars-CoV-2 proteins were obtained from the UniProt database (May 2021) and matched using the Andromeda^50^ algorithm at a peptide-spectrum match false discovery rate (FDR) of 0.05. Search parameters included a mass tolerance of 20 p.p.m. for the parent ion, 0.5 Da for the fragment ion, carbamidomethylation of cysteine residues (+57.021464 Da), variable N-terminal modification by acetylation (+42.010565 Da), and variable methionine oxidation (+15.994915 Da). Relative quantification was set as TMT 6-plex labels 126 to 131. The cleavage site specificity was set to Trypsin/P, with up to two missed cleavages allowed. Next, the evidence.txt and proteinGroups.txt were loaded into the R software (v4.0.2) for statistical analysis. The normalization and identification of differentially expressed proteins was performed using the MSstatsTMT package^51^. Multiple comparisons were corrected using the Benjamini-Hochberg approach.

### Leukocyte and lymphocyte isolation

For lymphocyte isolation, whole blood (2mL) was collected in 5mL polystyrene round-bottom heparinized vacutubes. To isolate lymphocytes by immunomagnetic negative selection, 100μL of Isolation Cocktail and 100μL of Rapid Spheres (EasySep™ Direct Human Total Lymphocytes Isolation Kit: 19655, StemCell Technologies) were added to 2 mL of whole blood. After mixing and 5min incubation at RT, the sample volumes were topped up to 2.5mL with 0.04% bovine serum albumin (BSA) in PBS. The diluted sample was incubated in the magnet without lid for 5min, at RT and negatively selected lymphocytes were decanted into a new 5 mL polystyrene tube. Except the addition of Isolation Cocktail, all steps were repeated once. The final lymphocyte cell suspension was transferred to a 15 mL polypropylene tube and a volume of 5mL 0.04% BSA in PBS was added to the sample. Lymphocytes were precipitated by centrifugation for 5 min at 2000rpm, supernatant was discarded, and cells were resuspended in 5 mL of 0.04% BSA in PBS. This last step was repeated once, and cells were then resuspended in 100 μL of PBS+0.04% BSA. Cell density was quantified with a hemacytometer, cell viability was assessed with Trypan Blue staining (T8154; Sigma Aldrich), and 7500 live lymphocytes were transferred to a sterile 1.5 mL microcentrifuge tube.

For leukocyte isolation, 1 mL of whole blood from heparin containing vacutubes was transferred to 5 mL polystyrene round-bottom tubes and 12μL of 0.5M EDTA was added. 2% FBS in PBS (1mL) and 50μL of EasySep RBC Depletion spheres (EasySep™ RBC Depletion Reagent: 18170, Stem Cell Technologies) were added to immunomagnetically deplete red blood cells. After 5 min of magnet incubation at RT, cell suspension containing leukocytes was decanted into a new 5mL polystyrene tube. To ensure complete removal of red blood cells, RBC depletion was repeated, and cell suspension containing leukocytes was decanted into a new 15mL polypropylene tube. Leukocytes were precipitated by centrifugation at 2000rpm for 5 min at 20°C and resuspended in 5mL of 0.04% BSA in PBS. This last step was repeated once, and leukocytes were resuspended in 2 mL of 0.04% BSA in PBS. Cell viability and cell density were assessed, and 7500 live leukocytes were transferred to the microcentrifuge tube containing the lymphocyte cell suspension. The volume of the cell suspension containing 7500 lymphocytes and 7500 leukocytes in a total of 50 μL of 0.04% BSA in PBS.

## Immunocytochemistry and immunohistochemistry

Isolated leukocyte and lymphocyte samples were fixed in 4% paraformaldahyde in PBS (0.2mM and pH7.4), and spun in a cytocentrifuge (8min at 300g) onto coated slides. Pathological lung sections (FFPE fixed and sectioned at 5um) were deparaffinized in Slide Brite (Fisher Scientific NC968653) and rehydrated. Slides were permeabilized and blocked with 10% normal donkey serum in PBS (with 0.5% triton X-100), primary antibodies (S100A8/9 Abcam ab22506; IFITM1 Abcam ab233545) were incubated at 4°C overnight, followed by incubation with donkey anti-rabbit-Alexa488 (Invitrogen A32790) or anti-mouse-Alexa555 (Invitrogen A31570) for 1h at room temperature (RT). Cytospun slides were sequentially stained with CD24 (Abcam ab202073) on the same slides for 1h at RT, followed by donkey anti-rabbit-Alexa647 (Invitrogen A31573). Imaging was done using a VS-120 slide scanner (Olympus) and high resolution image imaging was done using an SP8 spectral confocal microscope (Leica). Image processing was completed in Fiji ^52^.

### Single-cell RNA-Seq library construction, alignment, and quality control

A total of 15,000 single cells (containing an equal proportion of leukocytes and lymphocytes) were loaded for partitioning using 10X Genomics NextGEM Gel Bead emulsions (Version 3.1). All samples were processed as per manufacturer’s protocol (with both PCR amplification steps run 12X). Quality control of resulting libararies and quantification was performed using TapeStation D1000 ScreenTape assay (Agilent). Sequencing was performed using Illumina NovaSeq S2 and SP 100 cycle dual lane flow cells over multiple rounds to ensure each sample received approximately 32,000 reads per cell. Sequencing reads were aligned using CellRanger 3.1.0 pipeline^53^ to the standard pre-built GRCh38 reference genome. Samples that passed alignment QC were aggregated into single datasets using CellRanger aggr with between-sample normalization to ensure each sample received an equal number of mapped reads per cell. Aggregated non-dexamethasone-treated COVID-19 (n = 12) and bacterial ARDS (n = 9) samples recovered 1,872,659 cells that were sequenced to 38,410 post-normalization reads per cell. Likewise, aggregated COVID-19 samples with (n = 9) or without (n = 12) dexamethasone recovered 1,748,551 single cells sequenced to 51,415 post-normalization reads per cell. Aggregated healthy samples recovered 19,816 cells, including 1,912 post-QC neutrophils (n = 5).

### Single-cell RNA-Seq computational analyses and workflows

Filtered feature-barcode HDF5 matrices from aggregated datasets were imported into the R package Seurat v.3.9 for normalization, scaling, integration, multi-modal reference mapping, louvain clustering, dimensionality reduction, differential expression analysis, and visualization ^54^. Briefly, cells with abnormal transcriptional complexity (fewer than 500 UMIs, greater than 25,000 UMIs, or greater than 25% of mitochondrial reads) were considered artifacts and were removed from subsequent analysis. Since granulocytes have relatively low RNA content (due to high levels of RNases), QC thresholds were informed by ^8^ as they recently defined several rodent and human neutrophil subsets from scRNA-Seq samples. Cell identity was classified by mapping single cell profiles to the recently published PBMC single-cell joint RNA/CITE-Seq multi-omic reference ^55^.

### Annotation of neutrophil states

Since no published reference automates granulocyte annotations, neutrophil clusters were manually annotated by querying known markers (i.e. CSF3R, S100A8, S100A9, MMP8, MMP9, ELANE, MPO)^56^ and were corroborated using the R package SingleR^57^. Neutrophil states were defined by grouping unsupervised (louvain at default resolution) subclusters based on two overlapping criteria: scVelo-inferred neutrophil maturity, and 2. by corroborating gene expression and SCENIC-inferred GRN signatures with previous human and rodent neutrophil scRNA-Seq studies. Immature neutrophils were defined as CD24^+^ARG1^+^ELANE^+^MPO^+^ATF4^GRN-active^JDP2^GRN-active^ neutrophils ^7,8,47,58^that were reproducibly assigned as ‘root cells’ in scVelo-based latent time pseudo-ordering. IFN^active^ neutrophils were defined by preferential mRNA splicing (positive velocity) and expression of IFN-stimulated genes such as IFITM1/2, IFIT1/2/3, ISG15/20, and IFI6/27/44/44L ^6,44,59^. PG^active^ neutrophils were distinguished by preferential splicing of PTGS2/COX2 (as well as expression for prostaglandin transport LST1) ^44^ and included a subset that expressed high levels of IL1β decoy receptor IL1R2 ^33^. Lastly, IL7R^+^ neutrophils (a small but distinct subset that maybe of thymic origin ^60^ expressed high levels of ribosomal subunit genes (e.g. RPL5/7A/8/13/18/19/23/24/27/P0) that are highly reminiscent of ‘ribosomal^hi^-specific cluster 7’ identified previously ^47^..

### Statistical approach for comparing cell proportions

To test whether cell composition was changed due to infection type (COVID-19 versus Bacterial ARDS) or treatment group (dexamethasone versus non-dexamethasone), a generalized linear mixed-effects model was employed where infection type and treatment group were considered fixed and individual patients were considered random effect. Fitting was done with Laplace approximation using the ‘glmer’ function in the ‘lme4’ R package ^61^ and p-values were calculated using the R package ‘car’. Boxplots comparing cell type composition were generated using the ggplot2 package. Since a subset of patients sampled at t1 were discharged from ICU prior to t2 collection (non-random or non-ignorable missing data), we limit statistical comparisons to between group comparisons within one time point (e.g., COVID-19 72h vs Bacterial ARDS 72hr, dexamethasone-treated 72h vs non-dexamethasone-treated 72h) and do not estimate temporal differences across t1 and t2.

### Inferring cell communication networks

Differential cell-cell interaction networks were reconstructed using the Connectome R toolkit v0.2.2^62^ and CellChat v1.0.0 ^63^. Briefly, *DifferentialConnectome* queried Seurat R objects housing datasets integrated by infection type and dexamethasone status to define nodes and edges for downstream network analysis. Total number of interactions and interaction strengths were calculated using CellChat’s *compareInteractions* function. Differential edge list was passed through *CircosDiff* (a wrapper around the R package ‘circlize’) and CellChat’s *netVisual_chord_gene* to filter receptor-ligand edges and generate Circos plots.

### Consensus DEGs and perturbation scores

Differentially expressed genes (DEGs) were those with an average log fold change (FC) greater than 0.25 (p-adjusted < 0.05) as determined by Seurat’s Wilcoxon rank-sum test. Consensus stacked bars showing cumulative log fold changes (colored by individual sample contributions) were generated using *constructConsensus* function ^7^ for genes exhibiting reproducible changes across patients (>3 for 72-hour comparisons, > 2 for 7-day comparisons). Gene Set Enrichment analyses of consensus DEGs were performed using gProfiler’s g:GOSt (p-value cutoff <0.05). A cell state-specific ‘perturbation score’ was calculated to reflect the magnitude of response elicited by factoring in number and cumulative FC of consensus DEGs. Perturbation scores were visualized using Nebulosa-generated density plots ^64^.

### Constructing cellular trajectories using RNA velocity

Analysis of neutrophil trajectories was performed by realigning CellRanger count-generated BAMs with RNA velocity command-line tool ^20^ using the *run10x* command and human (GRCh38) annotations. The output loom files containing spliced and unspliced counts were combined to compare neutrophils in COVID-19 with Bacterial ARDS controls and dexamethasone-treated with non-treated COVID-19 patients. For both analyses, combined looms were imported into Seurat v.3.9 using the *ReadVelocity* function in SeuratWrappers v.0.2.0, normalized using *SCTransform* v.0.3.2 ^65^, reduced and projected onto a UMAP, and exported as a .h5 file using the *SaveH5Seurat* function. Counts stored in H5 files were imported, filtered, and normalized as recommended in the scVelo v.0.2.1 workflow ^19^. RNA velocities were estimated using stochastic and dynamical models. Since both models yielded comparable results, stochastic model was used as default for all subsequent analyses. Calculations stored in AnnData’s metadata were exported as CSVs and kernel density lines depicting Velocity-inferred latent time distribution were plotted with ggplot2 v.3.1.1.

### Gene Regulatory Network reconstruction

Single-cell regulatory network inference and clustering (SCENIC)^26^ was employed to infer regulatory interactions between transcription factors (TFs) and their targetome by calculating and pruning co-expression modules. Briefly, neutrophils were subsetted from scVelo-realigned Seurat object and processed using default and recommended parameters specified in SCENIC’s vignette (https://github.com/aertslab/SCENIC) using the hg19 RcisTarget reference. Regulon activity scores (in ‘3.4_regulonAUC.Rds’, an output of the SCENIC workflow) were added to scVelo object (using *CreateAssayObject* function) to jointly project trajectory and TF activity onto the same UMAP embeddings. Consensus stacked bars showing cumulative logFC of AUCell scores for each TF (colored by individual sample contributions) were generated by modifying the *constructConsensus* function^7^ for SCENIC assay. Targetome of TFs predicted as drivers of neutrophil states (stored in ‘2.6_regulons_asGeneSet.Rds’) was profiled using g:Profiler’s functional enrichment analysis and genes intersecting with the Interferon pathway were plotted using iRegulon (Cytoscape plugin)^66^.

### Comparing scRNA-Seq findings with published datasets

To test whether dexamethasone-suppressed neutrophil genes at t1 and t2 (Extended Data Table 4) predicted COVID-19 mortality, we repurposed methods described in ^33^ and employed whole blood bulk RNA-Seq datasets generated by ^34^ as a validation cohort of 103 samples (where 17 were fatal). Briefly, each of the 103 samples were scored by the aggregated expression of dexamethasone-suppressed neutrophil consensus genes at t1 and t2 using Seurat’s AddModuleScore(). Dexamethasone-suppressed module scores were used as the predictor variable and 28-day mortality was used as the response variable to construct an ROC curve using pROC’s roc() function. To infer bronchoalveolar neutrophil composition in severe and moderate COVID-19 ^11^and across bacterial pneumonia and COVID-19 ^32^, neutrophils (CSF3R^+^, S100A8^+^, S100A9^+^) captured in BALF scRNA-Seq datasets were projected onto our peripheral blood reference using mutual nearest neighbor anchoring (FindTransferAnchors) and identity transferring (TransferData and AddMetaData) strategy implemented in Seurat v4 ^54^.

### COVID Neutrophil Atlas

To enable intuitive exploration of single-cell datasets, a web portal (http://biernaskielab.ca/covid_neutrophil or http://biernaskielab.com/covid_neutrophil) was built using RShiny v1.1.0, shinyLP v.1.1.2, and shinythemes v.1.1.2 packages.

## Supporting information

Extended Figures 1-9 and Table Title

## Data availability

Single cell RNA-Seq datasets are available at NCBI GEO (which automatically makes SRA deposit) at the following accession: GSE157789. Single-cell datasets can be further explored on our companion portal at http://biernaskielab.ca/COVID_neutrophil or http://biernaskielab.com/COVID_neutrophil. Velocyto-generated LOOM files and processed R objects are available for reanalysis from: http://doi.org/10.6084/m9.figshare.14330795. Whole blood bulk RNA-Seq datasets employed as an independent validation cohort were downloaded from GSE157103. BALF scRNA-Seq datasets from severe and moderate COVID-19 were downloaded from GSE145926. Processed BALF scRNA-Seq objects from patients with bacterial pneumonia and COVID-19 (archived at GSE167118) were downloaded from authors’ archive: https://figshare.com/articles/dataset/_/13608734. Mass spectrometry datasets will be available via PRIDE Archive (http://www.ebi.ac.uk/pride/archive), it has been submitted (submission #: 1-20210702-114055) and is pending accessioning.

Proteomics data will be available at PRIDE (https://www.ebi.ac.uk/pride/), it has been submitted (submission #: 1-20210702-114055) and is pending accessioning.

## Code availability

All analyses were performed using publicly available software as described in the methods section. Raw scripts are available upon request.

## Supplementary Information

is available for this paper.

## Acknowledgements

This work was funded by a FastGrant from the Thistledown Foundation (JB and BY) and Calgary Firefighters Burn Treatment Society (JB). S Sinha received CIHR Vanier, Alberta Innovates, and Killam doctoral scholarships. E.L received an Alberta Children’s Hospital Research Institute postdoctoral fellowship. B.G.Y is a tier II Canada Research Chair in Pulmonary Immunology, Inflammation and Host Defence. We acknowledge the assistance of the nurse practitioners, Charissa Elton-Lacasse, Kirsten Deemer and Robert Ralph as well as the healthcare teams from the Calgary Adult ICU’s at South Health Campus, Rockyview General Hospital, Foothills Medical Center and Peter Lougheed Center. We thank Dr. Kirsten Fiest and the ICU study coordinators Cassidy Codan, Zdenka Slavikova and Olesya Dmitrieva. We acknowledge Dan Jones, Cathy Curr and the eCritical team (Alberta Health Services in Alberta, Canada) for their help in data acquisition and extraction via eCritical databases. Mortality predictions using dexamethasone-suppressed gene signatures were completed by repurposing computational workflows kindly shared by Aaron Wilk and Dr. Catherine Blish (Stanford University).

## Author contributions

SS performed scRNAseq analyses, figure preparation, and co-wrote the paper. NLR contributed to experimental design, performed scRNAseq experiments, figure preparation and co-wrote the paper. AJ, RA, and LC performed bioinformatics and created the online atlas. EL, RF and APN contributed to sample preparation and scRNAseq processing. MG and BM contributed to patient consent and sample collection. LGA and AD conducted proteomics and related analyses. AB provided clinical biospecimens. MJF provided serum cytokine assays. JB and BY conceived of all experiments, experimental design, wrote the paper and supervised all experiments.

**The authors have no competing interests.**

