## Extended Figures 1-9 and Table Title for "An Immune Cell Atlas Reveals Dynamic COVID-19 Specific Neutrophil Programming Amenable to Dexamethasone Therapy"

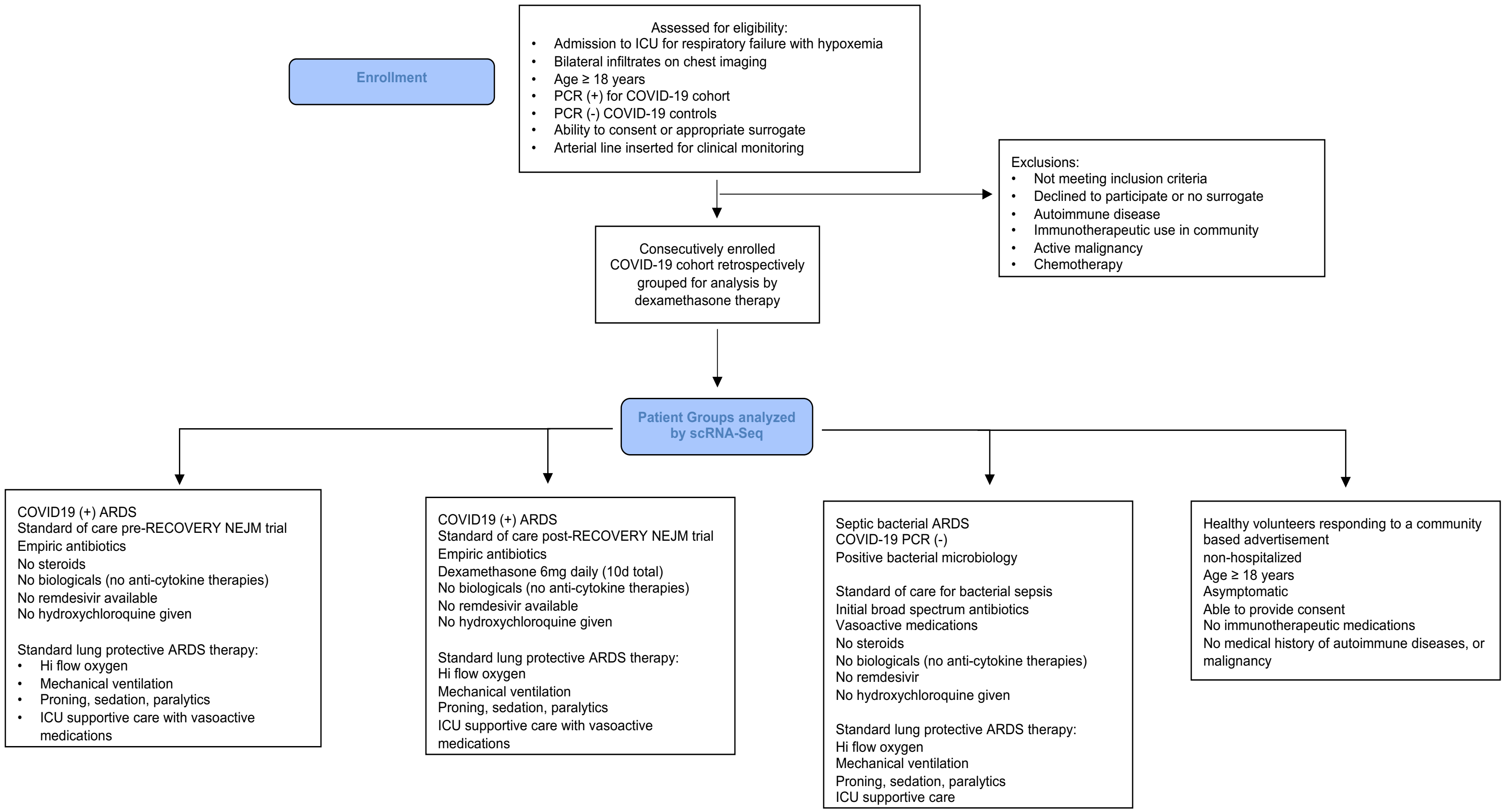


**Extended Data Figure 1. A modified CONsolidated Standards Of Reporting Trials (CONSORT) diagram showing trial groups in this study.**

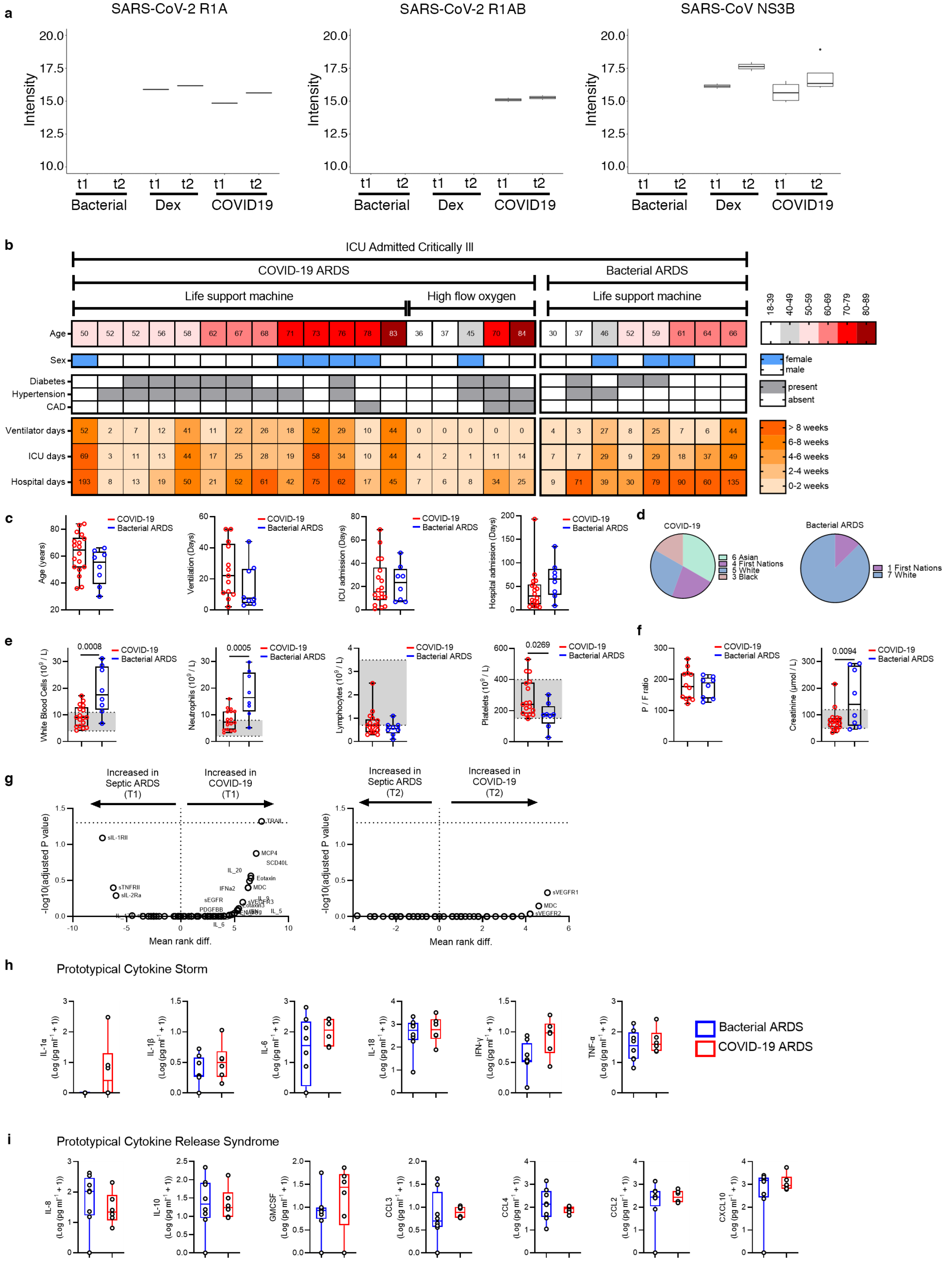
a
SARS-CoV-2 R1A
SARS-CoV-2 R1AB
SARS-CoV NS3B
Intensity
20.0
17.5
15.0
12.5
10.0
t1 t2 t1 t2 t1 t2
Bacterial Dex COVID19
b
ICU Admitted Critically Ill
COVID-19 ARDS
Bacterial ARDS
Life support machine
High flow oxygen
Life support machine
Age 50 52 52 56 58 62 67 68 71 73 76 78 83 36 37 45 70 84 | 30 37 46 52 59 61 64 66
18-39 40-49 50-59 60-69 70-79 80-89
Sex
female
male
Diabetes
Hypertension
CAD
present
absent
Ventilator days 52 2 7 12 41 11 22 26 18 52 29 10 44 0 0 0 0 0 | 4 3 27 8 25 7 6 44
ICU days 69 3 11 13 44 17 25 28 19 58 34 10 44 4 2 1 11 14 | 7 7 29 9 29 18 37 49
Hospital days 193 8 13 19 50 21 52 61 42 75 62 17 45 7 6 8 34 25 | 9 71 39 30 79 90 60 135
> 8 weeks
6-8 weeks
4-6 weeks
2-4 weeks
0-2 weeks
c
COVID-19
Bacterial ARDS
Age (years)
Ventilation (Days)
ICU admission (Days)
Hospital admission (Days)
d
COVID-19
6 Asian
4 First Nations
5 White
3 Black
Bacterial ARDS
1 First Nations
7 White
e
White Blood Cells (10^9 / L)
0.0008
Neutrophils (10^9 / L)
0.0005
Lymphocytes (10^9 / L)
Platelets (10^9 / L)
0.0269
f
P / F ratio
Creatinine (μmol / L)
0.0094
g
Increased in Septic ARDS (T1)
Increased in COVID-19 (T1)
Increased in Septic ARDS (T2)
Increased in COVID-19 (T2)
-log10(adjusted P value)
Mean rank diff.
TRAIL
sIL-1RII
MCP4
SCD40L
IL_20
Eotaxin
IFNa2
MDC
sTNFRII
sIL-2Ra
sEGFR
sVEGFR3
PDGFBB
IL_6
IL_5
sVEGFR1
MDC
sVEGFR2
h
Prototypical Cytokine Storm
IL-1α
IL-1β
IL-6
IL-18
IFN-γ
TNF-α
(Log (pg ml^-1 + 1))
Bacterial ARDS
COVID-19 ARDS
i
Prototypical Cytokine Release Syndrome
IL-8
IL-10
GMCSF
CCL3
CCL4
CCL2
CXCL10
(Log (pg ml^-1 + 1))

**Extended Data Figure 2. Clinical data of ICU admitted COVID-19 and septic bacterial ARDS**
a) Shotgun proteomics assessment using tandem Mass Spectroscopy with a targeted search run for known SARS-CoV-2 proteins R1A and R1AB, and SARS-CoV protein NS3B are displayed for all patient cohorts. b) Summary of individual information of ICU admitted patients with established COVID-19 or a diagnosis of bacterial ARDS due to sepsis. Age, sex, comorbidities and lengths of stay are displayed. c) Aggregated cohort clinical data and d) racial backgrounds. e) Clinical cell counts from peripheral blood taken on t1; shaded areas show local lab normal values. f) PaO2/FiO2 ratio (P/F) and creatinine at t1. g) multiple comparison analysis of all serum cytokines assessed at t1 and t2 are shown as volcano plots. Serum cytokine determination of prototypical mediators involved in h) cytokine storm and i) cytokine release syndrome graphed in Log transformation taken at t1.

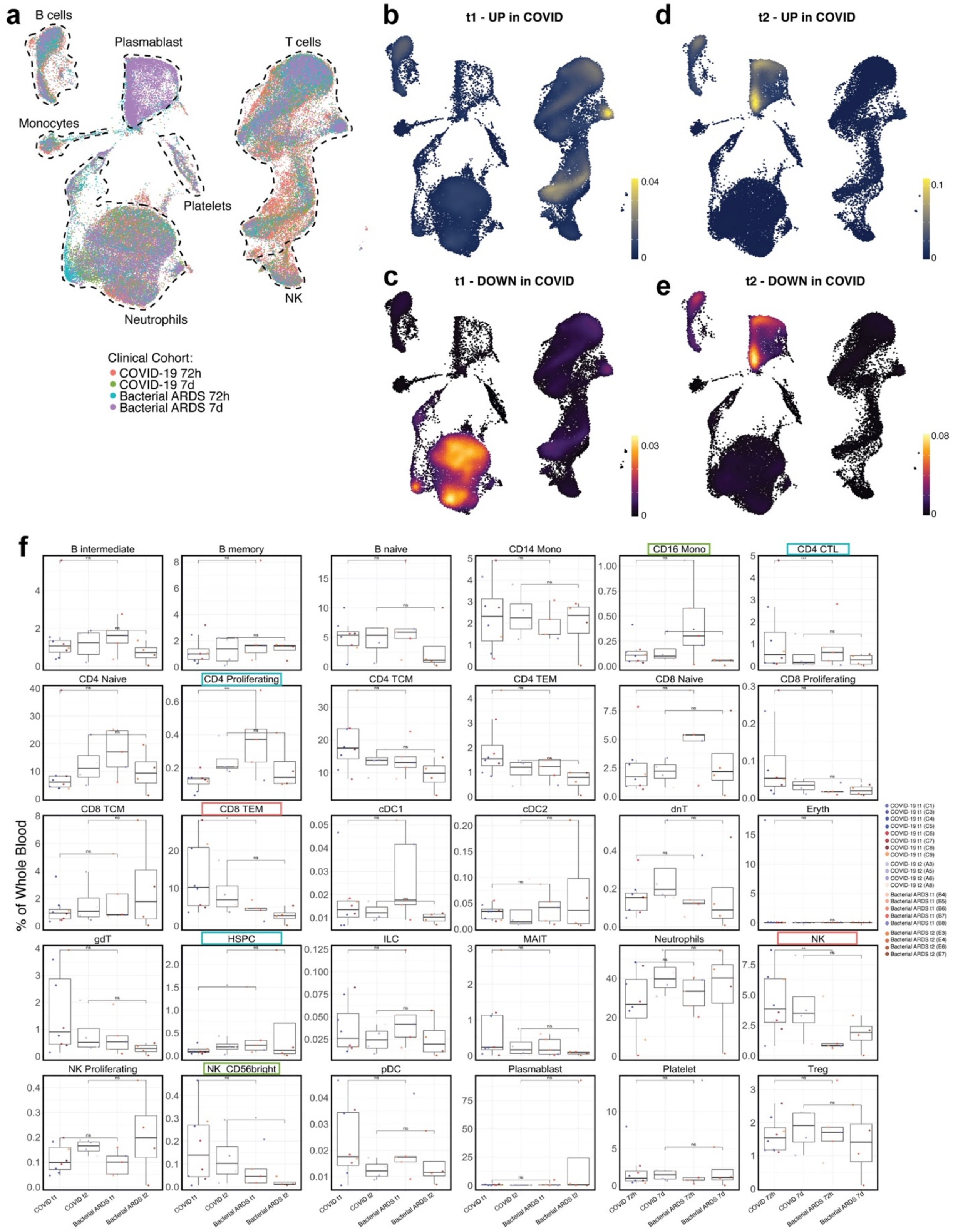


**Extended Data Figure 3. COVID-19 elicits distinct innate and adaptive immune responses compared to bacterial ARDS.** a. UMAP projection of 86,935 whole blood cells from 21 patient samples, coloured by clinical cohort. b-e. Kernel density estimates depicting magnitude of response elicited by immune cell subsets in COVID-19 t1 (b-c) and t2 (d-e) calculated by summing consensus DEG fold changes for each cell subset shown in Panel B. Consensus DEGs upregulated in COVID-19 are plotted on cividis spectrum (yellow = higher expression) whereas downregulated DEGs are plotted on inferno spectrum (yellow/orange = lower expression). f. Boxplots showing percentage of each cell type in each patient sample grouped by clinical cohort and coloured by donor ID. The x axes correspond to the clinical cohort of each patient. Biologically independent samples for COVID-19 at t1 ($n = 8$), COVID-19 at t2 ($n = 4$), bacterial ARDS at t1 ($n = 5$), and bacterial ARDS at t2 ($n = 4$). Significance of effects were estimated using logistic regression models as comparisons included fixed (COVID-19 vs Bacterial ARDS) and random (per patient) effects. All effects with $p < 0.05$ are indicated. *$p < 0.05$, **$p < 0.01$, ***$p < 0.001$.

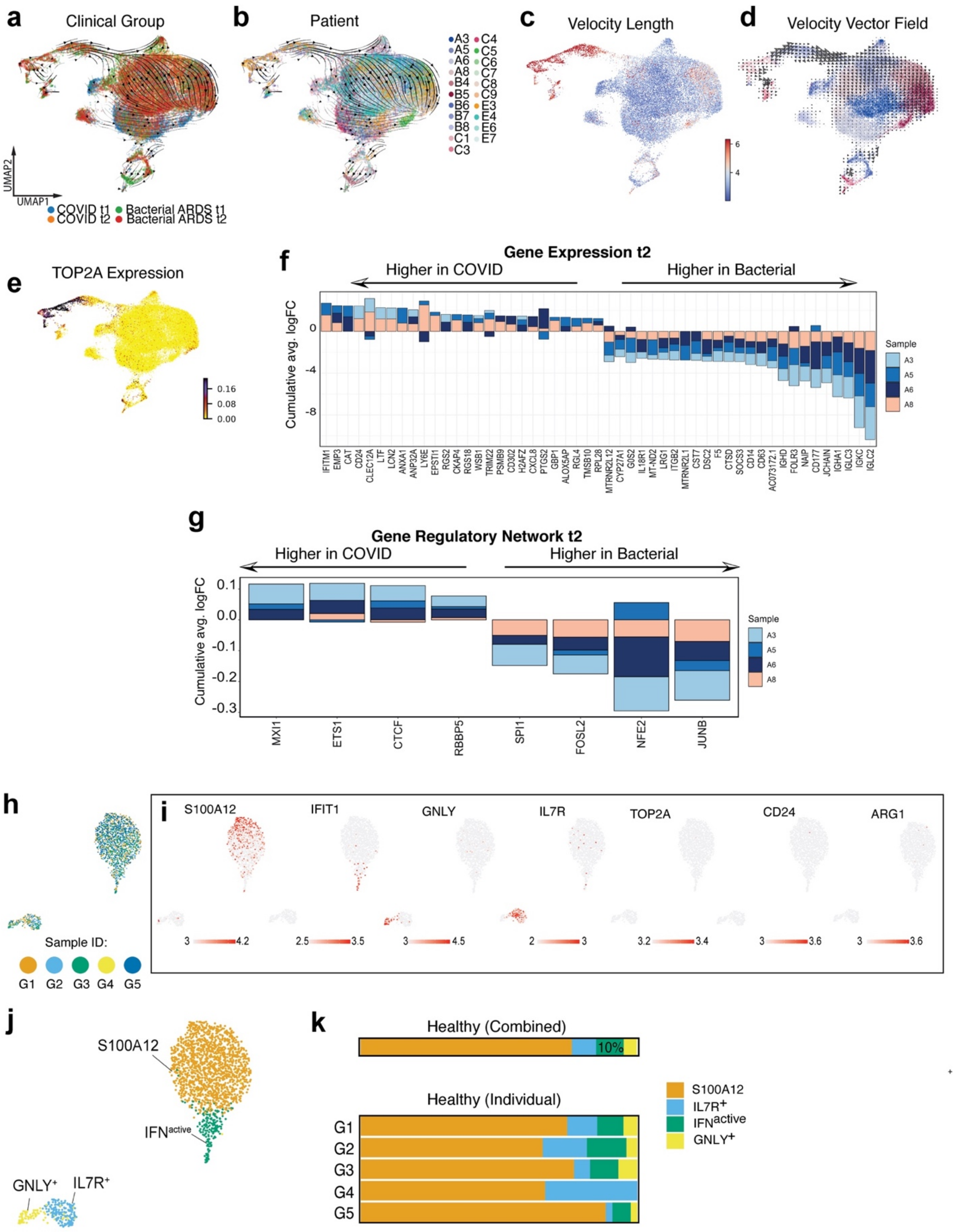
a
Clinical Group
UMAP2
UMAP1
COVID t1
COVID t2
Bacterial ARDS t1
Bacterial ARDS t2
b
Patient
A3
A5
A6
A8
B4
B5
B6
B7
B8
C1
C3
C4
C5
C6
C7
C8
C9
E3
E4
E6
E7
c
Velocity Length
6
4
d
Velocity Vector Field
e
TOP2A Expression
0.16
0.08
0.00
f
Gene Expression t2
Higher in COVID
Higher in Bacterial
Cumulative avg. logFC
0
-4
-8
Sample
A3
A5
A6
A8
IFITM1 EMP3 CAT CD24 CLEC12A LTF LCN2 ANXA1 ANP32A LY6E EPSTI1 RGS2 CKAP4 RGS18 WSB1 TRIM22 PSMB9 CD302 H2AFZ CXCL8 PTGS2 GBP1 ALOX5AP RGL4 TMSB10 RPL28 MTRNR2L12 CYP27A1 G0S2 IL18R1 MT-ND2 LRG1 ITGB2 MTRNR2L1 CST7 DSC2 F5 CTSD SOCS3 CD14 CD63 AC073172.1 IGHD FOLR3 NAIP CD177 JCHAIN IGHA1 IGLC3 IGKC IGLC2
g
Gene Regulatory Network t2
Higher in COVID
Higher in Bacterial
Cumulative avg. logFC
0.1
0.0
-0.1
-0.2
-0.3
Sample
A3
A5
A6
A8
MXI1
ETS1
CTCF
RBBP5
SPI1
FOSL2
NFE2
JUNB
h
Sample ID:
G1
G2
G3
G4
G5
i
S100A12
IFIT1
GNLY
IL7R
TOP2A
CD24
ARG1
3
4.2
2.5
3.5
3
4.5
2
3
3.2
3.4
3
3.6
3
3.6
j
S100A12
IFN^active
GNLY⁺
IL7R⁺
k
Healthy (Combined)
10%
Healthy (Individual)
G1
G2
G3
G4
G5
S100A12
IL7R⁺
IFN^active
GNLY⁺

**Extended Data Figure 4. COVID-19 infection reprograms neutrophil maturation by driving expansion of ISG- and PG-expressing subsets. a-d**. UMAPs plotting velocity analysis of 29,653 subclustered neutrophils undergoing state transitions, coloured by clinical cohort (a), patient ID (b), magnitude velocity vector length reflecting the difference between expected versus recovered unspliced counts (c), and neutrophil louvain clusters overlaid with velocity vector fields (d). **e.** Expression of immature neutrophil marker TOP2A. **f-g.** Consensus plot of differentially expressed genes (f) and SCENIC-inferred transcription factors (g) upregulated (positive logFC) or suppressed (negative logFC) in neutrophils from at least 2 of 4 patients with COVID-19 relative to bacterial ARDS at t2. **h.** UMAP of neutrophils from healthy donors (n = 1,912 cells) colored by donor of origin. **i.** Expression of state-defining neutrophil markers. **j.** UMAP of neutrophils from healthy donors colored by neutrophil states. k. Neutrophil state composition in healthy donors, combined across all donors or separated by individual donor ID.

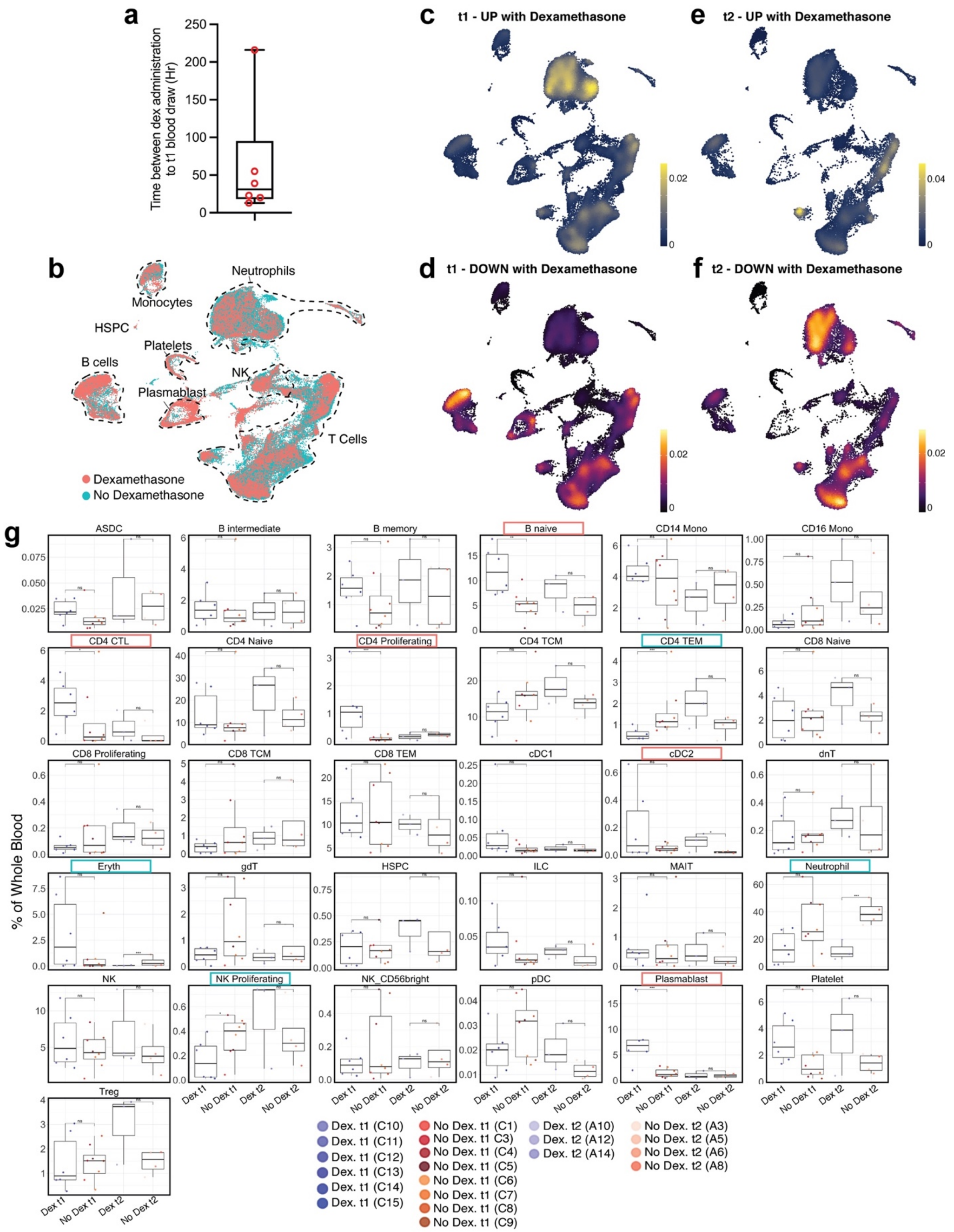


**Extended Data Figure 5. Immunosuppressive effects of dexamethasone are mediated through neutrophils and multiple adaptive immune cell subsets.** a. Bar plot shows distribution of time interval (in hours) between dexamethasone administration to first blood draw at t1. Data are mean ± SEM. b. UMAP projection of 80,994 whole blood cells from 21 patient samples, coloured by treatment groups. c-f. Kernel density estimates depicting magnitude of response elicited by immune cell subsets following dexamethasone treatment at 72 hours post-ICU (c-d) and 7 days post-ICU (e-f) calculated by summing consensus DEG fold changes for each cell subset shown in Panel B. Consensus DEGs upregulated following dexamethasone treatment are plotted on cividis spectrum whereas downregulated DEGs are plotted on inferno spectrum. g. Boxplots showing percentage of each cell type in each patient sample grouped by treatment and coloured by donor ID. The x axes correspond to four treatment groups. $n = 6$, $n = 8$, $n = 3$, and $n = 4$ biologically independent samples from dexamethasone-treated t1, no dexamethasone t1, dexamethasone-treated t2, no dexamethasone t2, respectively. Significance of effects were estimated using logistic regression models as comparisons included fixed (dexamethasone vs no dexamethasone) and random (per patient) effects. All effects with $p < 0.05$ are indicated. *$p < 0.05$, **$p < 0.01$, ***$p < 0.001$.

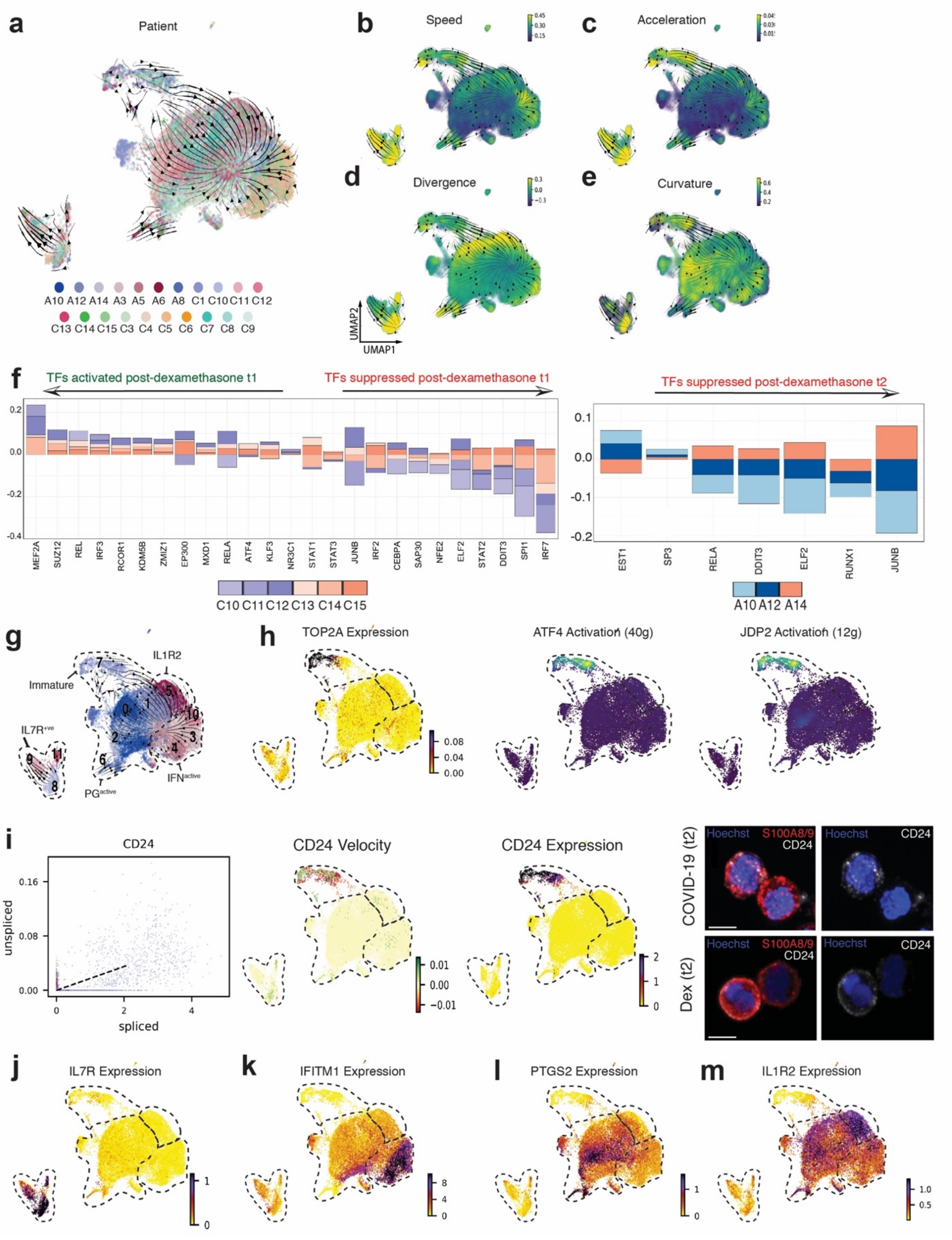


**Extended Data Figure 6. Distinct neutrophil states and their response to dexamethasone. a.** UMAP projection of subclustered neutrophils from 21 patient samples, coloured by individual patient ID. **b-e**. Cell speed (length of velocity vectors; b), acceleration (subspaces where velocity undergoes dramatic changes in either in magnitude or direction; c), divergence (outward flux indicating the extent to which a point behaves like a source; d) and curvature (hotspots of abrupt vector field change; e). **f**. Differentially activated consensus TFs upregulated (positive logFC) or suppressed (negative logFC) post-dexamethasone in at least 3 of 6 patients at t1, and in at least 2 of 3 patients at t2. **g-j**. Neutrophil states (g) can be distinguished by expression of proliferative marker TOP2A and activation of immaturity-associated TFs ATF4 and JDP2 (h), CD24 splicing kinetics, velocity, expression and immunocytochemistry (i), IL7R (j), Interferon-stimulated genes such as IFITM1 (k), genes involved in prostaglandin synthesis such as PTGS2 (l), and IL1R2 (m).

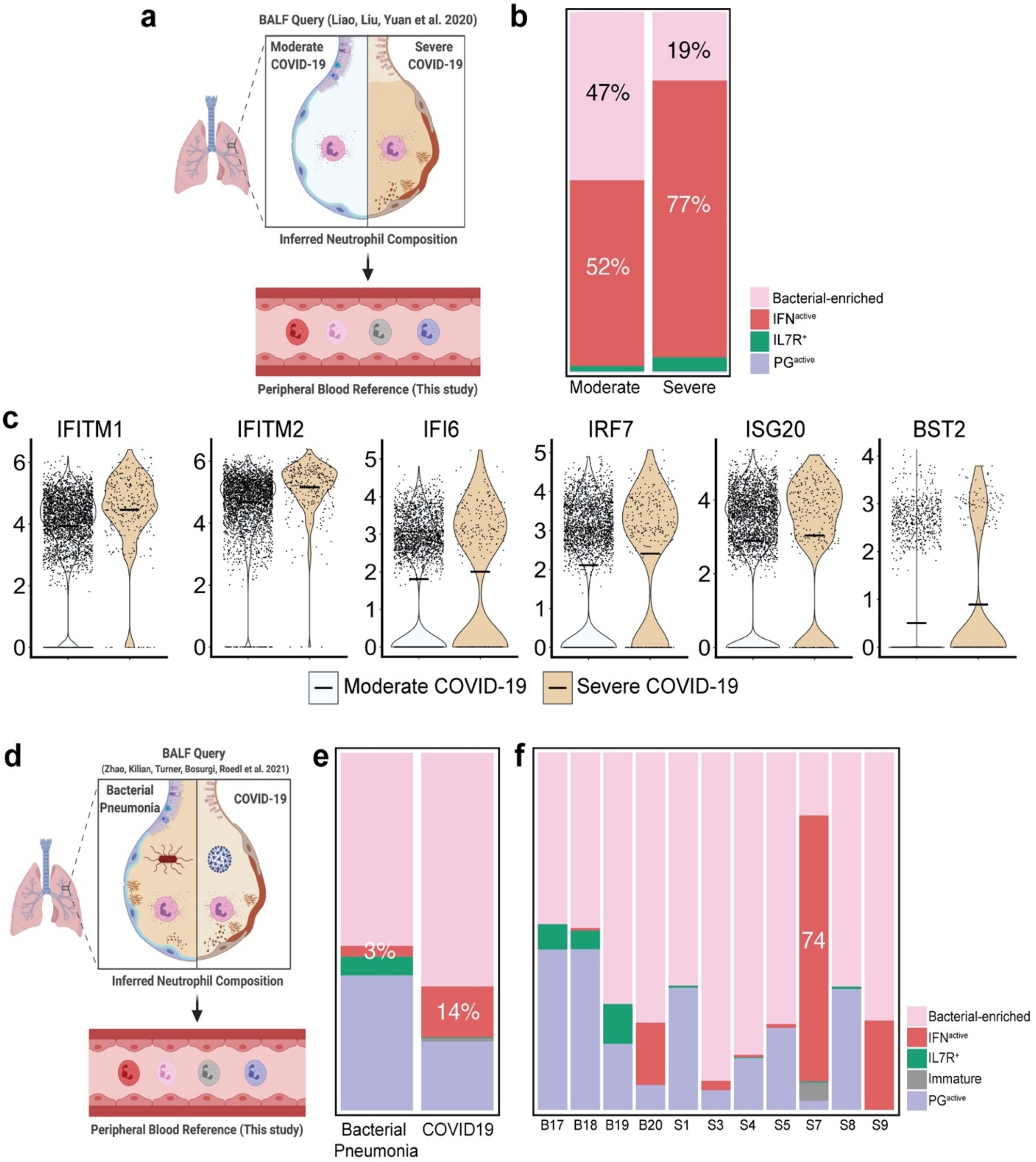

**Extended Data Figure 7. Inferred neutrophil composition in bronchoalveolar microenvironments.** **a.** Strategy for inferring BALF neutrophil composition in severe and moderate COVID-19 by reference-projecting to neutrophil states in peripheral blood. **b.** Proportion of neutrophil states in bronchoalveolar microenvironment in severe and moderate COVID-19. **c.** Expression of Type 1 IFN genes in neutrophils across severe and moderate COVID-19 patients. **d.** Strategy for inferring BALF neutrophil composition in bacterial pneumonia versus COVID-19. **e-f.** Proportion of neutrophil states in bronchoalveolar microenvironment, separated by bacterial pneumonia and COVID-19 (e) and individual donors (f).

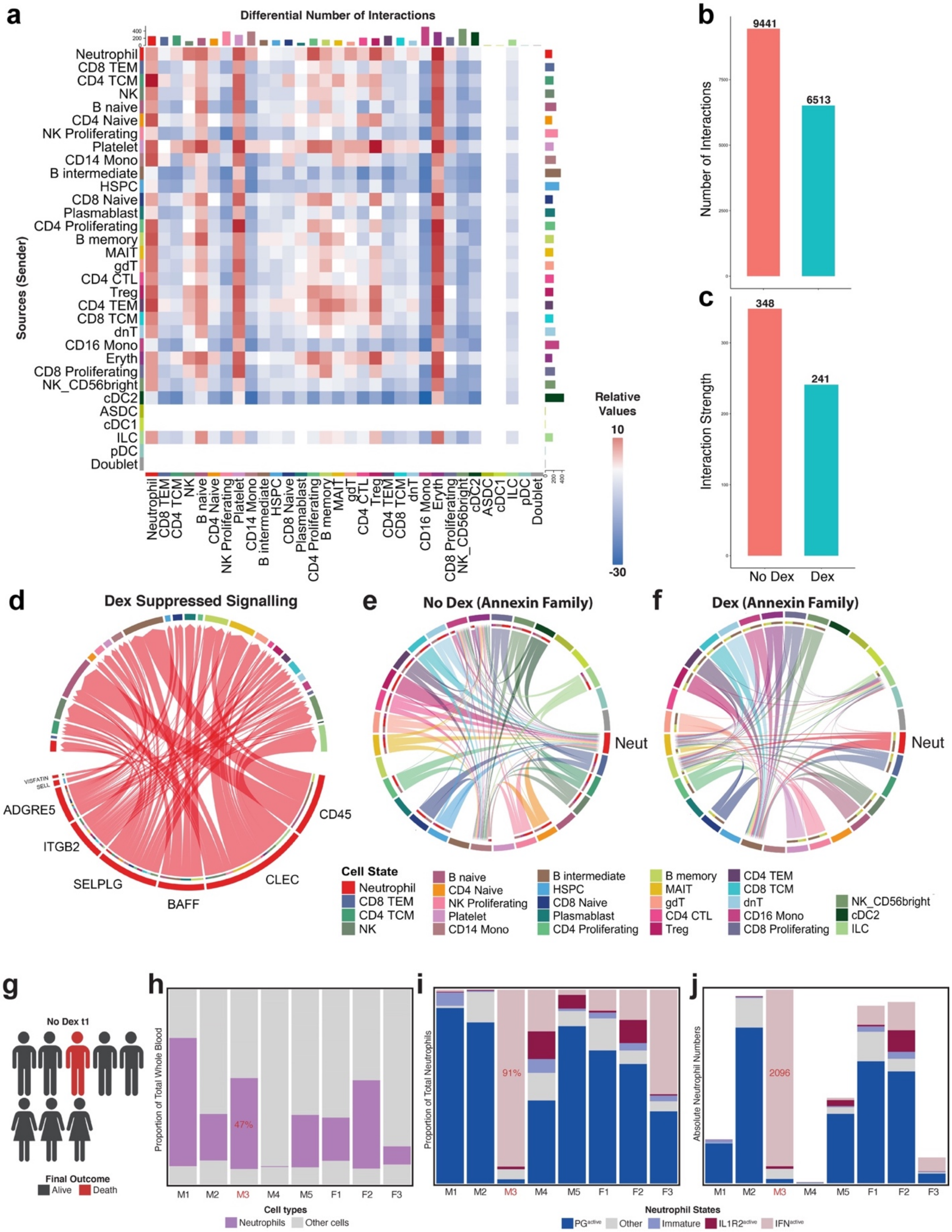


**Extended Data Figure 8. Dexamethasone alters global signaling topology and increased proportion of IFN$^{active}$ neutrophils are associated with mortality. a.** Interaction heatmap summarizing differential number of incoming (top bar plot) and outgoing (right bar plot) cell-to-cell interactions following dexamethasone treatment. **b-c.** Global summary of number (b) and strength (c) of all interactions different immune cell types with and without dexamethasone. **d.** Neutrophil-driven signaling pathways supressed post-dexamethasone, identified using CellChat (MHC-I signaling filtered out). **e-f.** Unfiltered topology of annexin signaling without (e) and with dexamethasone (f) treatment. **g.** Schematic depicting outcomes in non-dexamethasone treated COVID-19 patients. Male 3 (M3) succumbed to disease. **h-i.** Proportion of neutrophils (h) and neutrophil states (i) in whole blood samples from individual donors (5 males, 3 females) at t1. **j.** Raw neutrophil state counts from the same 8 donors at t1.

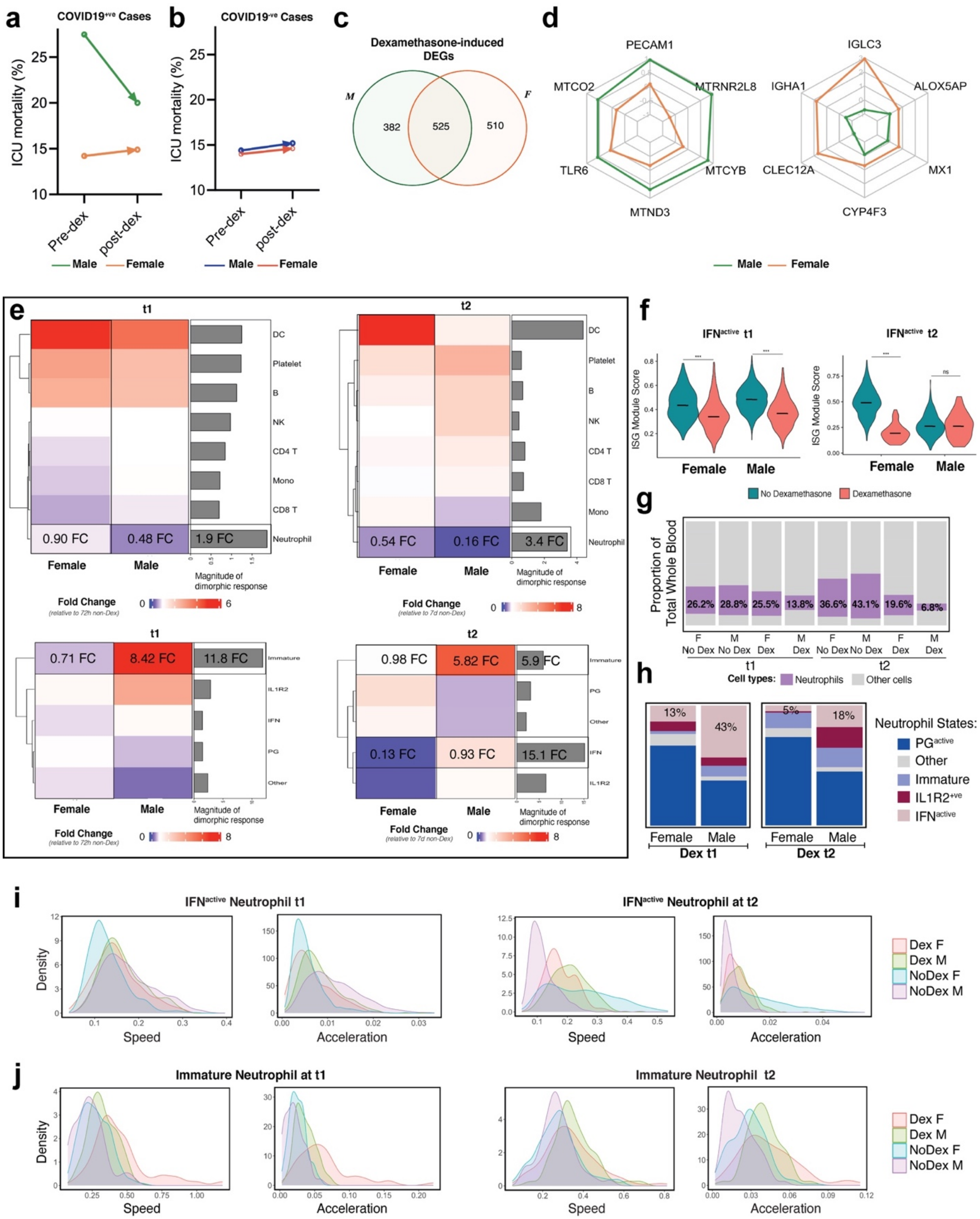


**Extended Data Figure 9. Dexamethasone attenuates neutrophil response in a sexually dimorphic fashion. a-b.** ICU mortality rates of sex-separated patients with (a) or without COVID-19 (b) comparing pre-dexamethasone (January 2020 till May 31st, 2020) and post-dexamethasone (June 1st, 2020, till May 31st, 2021) standard of care time periods. **c.** Number of genes that are uniquely or jointly regulated with dexamethasone between males and females. **d.** Differential magnitude or direction of regulation within dexamethasone-induced DEGs jointly regulated by both sexes. **e.** Heatmap depicting dexamethasone-induced shifts in cellular composition at t1 and t2 and accompanying bar plots showing magnitude of divergence between male and female response**.** Dexamethasone-induced shifts in neutrophil state composition at t1 and t2 along with magnitude of divergence between male and female response. **f.** Module score of ISG signatures in ISG-active neutrophils across sex and dexamethasone treatment at t1 and t2. Statistical significance was assessed using an ANOVA test followed by bonferroni-corrected pair-wise t-tests. * p-value < 0.05; ** p-value < 0.01; *** p-value < 0.001; ns p-value > 0.05. Center line indicates median data point. **g**. Comparison of proportion of neutrophils in whole blood samples from sex-separated cohorts. **h**. Comparison of neutrophil composition across sex in dexamethasone-treated patients at 72 hours and 7 days post-ICU admission. **i-j**. Histograms depicting cell speed (length of velocity vectors) and acceleration (subspaces where velocity undergoes dramatic changes in magnitude or direction) in IFN-active (i) and immature (j) neutrophils, separated by sex and dexamethasone treatment for both t1 and t2.

**Extended Data Table 1 –** Clinical and demographic characteristics of all analyzed donors in COVID-19 versus bacterial ARDS and dexamethasone versus non-dexamethasone COVID-19 controls at t1 and t2 post-ICU admission.

**Extended Data Table 2 –** Targeted list of shotgun proteomics comparing a) COVID-19 and bacterial ARDS serum, or b) COVID-19 and dexamethasone treated patients. Tab 1 lists proteins that are significantly different (adjusted p-value <0.05) and tab 2 lists the proteins that fall outside of the interquartile range (the IQR is listed for each comparison).

**Extended Data Table 3 –** Consensus DEGs in each major cell type in patients with COVID-19 relative to bacterial ARDS 72 hours and 7 days post-ICU admission. Reported are average LogFC change of DEGs comparing patients with COVID-19 at t1 (columns C1, C3-C9) and t2 (columns A3, A5, A6, A8) to bacterial ARDS controls.

**Extended Data Table 4 –** Consensus DEGs in each major cell type in COVID-19 patients treated with dexamethasone relative to non-dexamethasone COVID-19 controls t1 and t2 post-ICU admission. Reported are average LogFC change of DEGs comparing patients with dexamethasone at t1 (columns C10-15) and t2 (columns A10, A12, A14) to non-dexamethasone treated COVID-19 controls.

**Extended Data Table 5 –** Sexually dimorphic neutrophil DEGs in COVID-19 patients treated with dexamethasone relative to non-dexamethasone COVID-19 controls at t1 post-ICU admission. Reported are neutrophil transcriptome modulated by dexamethasone in both sexes (columns B-G), in males alone (columns I-N), and in females alone (columns P-S).

**Extended Data Video 1 –** Dynamo-reconstructed neutrophil vector field topology (left) and animation depicting fate commitments towards COVID-19 and Bacterial ARDS-enriched states (right).
